# Microfluidics-free single-cell genomics with templated emulsification

**DOI:** 10.1101/2022.06.10.495582

**Authors:** Iain C. Clark, Kristina M. Fontanez, Robert H. Meltzer, Yi Xue, Corey Hayford, Aaron May-Zhang, Chris D’Amato, Ahmad Osman, Jesse Q. Zhang, Pabodha Hettige, Jacob S.A. Ishibashi, Cyrille L. Delley, Daniel W. Weisgerber, Joseph M. Replogle, Marco Jost, Kiet T. Phong, Vanessa E. Kennedy, Cheryl A. C. Peretz, Esther A. Kim, Siyou Song, William Karlon, Jonathan S. Weissman, Catherine C. Smith, Zev J. Gartner, Adam R. Abate

**Author notes:** Corresponding author: Adam R. Abate.

## Abstract

Single-cell RNA sequencing is now a standard method used to reveal the molecular details of cellular heterogeneity, but current approaches have limitations on speed, scale, and ease of use that stem from the complex microfluidic devices or fluid handling steps required for sample processing. We, therefore, developed a method that does not require specialized microfluidic devices, expertise, or hardware. Our approach is based on particle-templated emulsification, which allows single-cell encapsulation and barcoding of cDNA in uniform droplet emulsions with only a vortexer. PIP-seq accommodates a wide range of emulsification formats, including microwell plates and large-volume conical tubes, enabling thousands of samples or millions of cells to be processed in minutes. We demonstrate that PIP-seq produces high-purity transcriptomes in mouse-human mixing studies, is compatible with multi-omics measurements, and can accurately characterize cell types in human breast tissue when compared to a commercial microfluidic platform. Single-cell transcriptional profiling of mixed phenotype acute leukemia using PIP-seq revealed the emergence of heterogeneity within chemotherapy-resistant cell subsets that were hidden by standard immunophenotyping. PIP-seq is a simple, flexible, and scalable next-generation workflow that extends single-cell sequencing to new applications, including screening, diagnostics, and disease monitoring.

## Introduction

Single-cell RNA sequencing (scRNA-seq) is an essential technology in the biological sciences because it reveals how the properties of tissues arise from the transcriptional states of numerous interacting cells. Defining the gene expression signatures of individual cells allows cell type classification, the discovery of unique cell states during development and disease, and the prediction of regulatory mechanisms that control these states. As a result, bulk sequencing is being rapidly replaced by single-cell methods. The first single-cell approaches isolated cells and prepared them individually for sequencing^1–4^. While improvements in molecular biology increased data quality^5,6^, the requisite isolation and processing of separate cells ultimately limited throughput. Implementation of valve-based microfluidics reduced hands-on time^7^, but failed to significantly increase cell number, and thus could not capture the heterogeneity intrinsic to most tissues. Advances in high throughput droplet microfluidic barcoding have expanded single-cell sequencing to tens of thousands of cells^8,9^ and fueled biological discovery, but require expensive instruments located in core facilities and therefore remain inaccessible to many labs. Methods for direct combinatorial indexing of cells^10,11^, the use of nanowell arrays ^12^, or sample multiplexing^13,14^ have overcome some limitations of microfluidics, but no current method simultaneously accommodates both low (10) and high (>10^6^) cell numbers, can be applied to hundreds of independent samples and can be rapidly implemented without custom equipment.

The scalability of single-cell methods is important for many applications, including tissue atlas projects^15–18^, million cell perturbation experiments^19^, drug development pipelines^20^, and developmental studies^21^. Droplet microfluidics has an intrinsic disadvantage at high cell numbers due to the upper limit on drop generation speed. At high fluid velocities, droplet generation becomes uncontrolled, resulting in polydispersed emulsions and poor bead loading that reduces single-cell data quality^22,23^. Therefore, to sequence millions of cells requires long run times, parallel droplet generators with complex designs that are prone to clogging, or implementation of additional barcoding steps before encapsulation^24^. More generally, droplet microfluidics relies on an expensive instrument usually located in a core facility, which necessitates sample transport or fixation that can alter RNA profiles. Centralized processing also reduces access to many labs and does not fit experiments that need rapid or point-of-collection sample handling, like remote fieldwork or studies using infectious samples requiring biosafety precautions^12,25^.

Much effort has thus gone into developing microfluidic-free single-cell methods. Split-pool ligation^10,11^ and tagmentation^26,27^ perform direct combinatorial barcoding of bulk suspensions, and significantly increase cell number; however, these laborious workflows require enormous numbers of pipetting operations and are poorly suited for low cell inputs. Moreover, while scalable, these methods require substantial expertise^28^, and broad adoption of split-pool barcoding will likely require robotic automation in a centralized facility. Alternatively, methods based on nanowells prioritize simplicity and cost-effectiveness^12,25^. No microfluidics are required and wells are loaded by sedimentation, providing an instrument-free and point-of-use solution. However, nanowell array chips do not efficiently scale in cell or sample number; the planar arrays capture cells on a 2D surface and, thus, cannot compete with emulsions or combinatorial indexing utilizing a 3D volume that easily scales to millions of cells. Moreover, unless combined with multiplexing^13,14^, nanowell chips are poorly suited for processing many separate samples because they require one array per sample, and thus hundreds of arrays for hundreds of samples. To advance the field of single-cell genomics, next-generation technologies must simultaneously innovate on speed, scale, and ease of use. An ideal system would be compatible with the barcoding of separate samples in well plates, accommodate orders-of-magnitude differences in cell number, be completed in minutes, and be easy to run at the bench or in the field without specialized instrumentation. An approach that achieved these innovations simultaneously would significantly enhance the usefulness of single-cell genomics, impact basic and clinical research, and facilitate new diagnostics.

Here we describe a flexible, scalable, and instrument-free scRNA-seq method based on rapid templated emulsification of cells and barcoded hydrogel templates without microfluidics^29^. In contrast to microfluidic emulsification, in which droplets are created sequentially and thus their number scales with instrument run time, templated emulsification generates monodispersed droplets in parallel by bulk self-assembly and thus the number of droplets (and cells that can be barcoded) scales only with container volume. The result is an extremely scalable, user-friendly scRNA-seq method that we call PIP-seq (**P**article-templated **I**nstant **P**artition-seq). Templated emulsification produces drops that are equivalent to those generated with microfluidics and compatible with the latest innovations in multi-omic measurements. Here, we show that PIP-seq generates accurate single-cell gene expression profiles from human tissues and is compatible with multi-modal measurements of RNA and sgRNA (CROP-seq), or RNA and protein (CITE-seq). Finally, we demonstrate the use of PIP-seq to monitor the response of patients with mixed phenotype acute leukemia (MPAL) to chemotherapy, revealing heterogeneity within cells with similar immunophenotypes. In summary, PIP-seq fills an unmet technical need by improving the speed, scalability, and ease of use of single-cell sequencing.

## Results

### Overview of the technology

PIP-seq uses particle templating to compartmentalize cells, barcoded hydrogel templates, and lysis reagents in monodispersed water-in-oil droplets (**Fig. 1a**). Rapid emulsification with a standard vortexer allows cells to be encapsulated at the bench or point of collection in minutes. The cells are lysed by increasing the temperature to 65°C, which activates Proteinase K, releasing cellular mRNA that is captured on polyacrylamide beads decorated with barcoded polyT sequences (**Fig. 1b**). PIP-seq emulsions can be stored for days at 0°C without change in data quality **(Extended data Fig. 1)**, allowing samples to be banked for future processing. Upon resuming, oil is removed, beads are transferred into a reverse transcription buffer, and full-length cDNA is synthesized, amplified, and prepared for sequencing (**Fig. 1c-d**).

**Figure 1.**
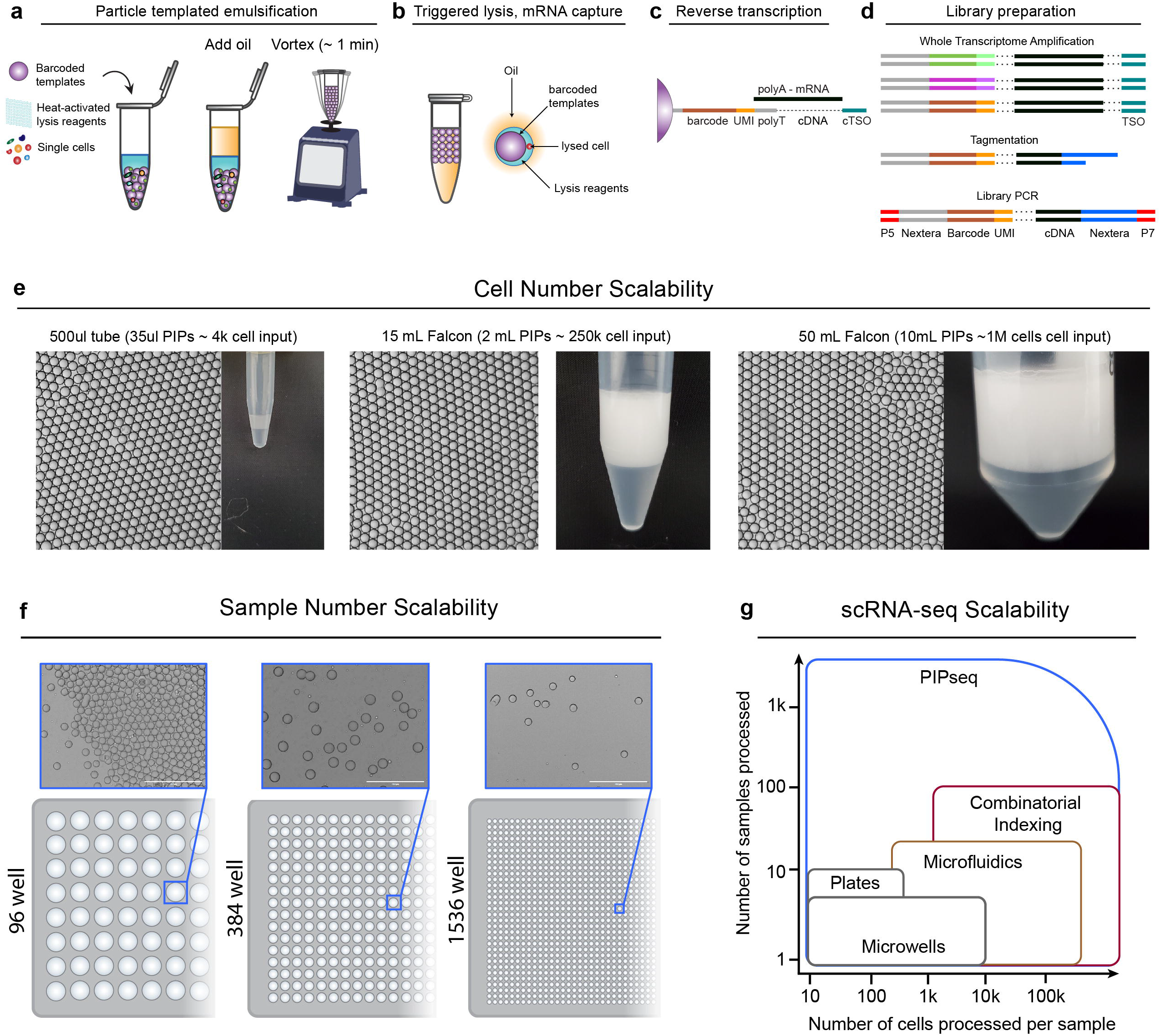
Rapid and scalable templated emulsification for single-cell genomics. **(a-d)** PIP-seq enables the encapsulation, lysis, and barcoding of single cells. **(a)** Schematic of the emulsification process. Barcoded particle templates, cells, and lysis reagents are combined with oil and vortexed to generate monodispersed droplets. **(b)** Heat activation of Proteinase K results in lysis and release of mRNA that is captured on bead-bound barcoded polyT oligonucleotides. **(c)** Oil removal is followed by bulk reverse transcription of mRNA into cDNA. **(d)** Barcoded whole transcriptome amplified cDNA is prepared for Illumina sequencing. **(e-g)** Efficient single bead, single drop encapsulation at scale. **(e)** Particle templated emulsification in different sized tubes (1.5 ml, 15 ml, 50 ml) produces monodispersed emulsions capable of barcoding orders of magnitude different cell numbers. **(f)** PIP-seq is compatible with plate-based emulsification, including 96-, 384- and 1536-well plate formats. **(g)** The estimated ability of different technologies to easily scale with respect to cell and sample number.

A unique and valuable feature of PIP-seq is that cell encapsulation in droplets is performed in parallel using bead size to control droplet volume. In contrast to microfluidics, the number of droplets scales with total container volume, not emulsification time. For example, at a 6% collision rate that includes cell doublets and barcode reuse, we estimate that 3.5k cells can be barcoded with 35 μL of barcoded hydrogel templates in a 500 μL tube, 225k cells can be barcoded with 2 mL of barcoded hydrogel templates in a 15mL conical tube, and 1M cells can be barcoded with 10mL of barcoded hydrogel templates in a 50mL conical tube (**Fig. 1e**). Regardless the tube size, only two minutes of vortexing is required for cell capture. PIP-seq is equally scalable to large sample numbers. Encapsulation can be performed directly in 96, 384, or 1536 well plates (**Fig. 1f, Extended data Fig. 2**), greatly simplifying experiments testing hundreds of different conditions and streamlining integration with robotic handling systems. Thus, compared to current single-cell RNA-sequencing technologies, PIP-seq has the greatest flexibility to cover combinations of cell and sample numbers (**Fig. 1g**).

### Single-cell RNA-sequencing with particle-templated emulsification

High-throughput single-cell sequencing requires efficient cell lysis and reverse transcription of mRNA using barcoded primers. In the absence of microfluidics, barcoded hydrogel templates, cells, and lysis reagents must be combined before emulsification. To prevent cell lysis before compartmentalization, we use Proteinase K (PK), a protease that has minimal activity at 4°C but can be activated at higher temperatures. After emulsification, the sample is heated to efficiently lyse cells. To illustrate this process, we stained cells with calcein, performed templated emulsification at 4°C with PK, and imaged the droplets before and after thermal activation. Intact cells appeared as compact puncta before lysis, but rapidly released calcein into the bulk of the droplets after the temperature was increased (**Fig. 2a, Extended data Fig. 2a,b**). Thus, cells can be mixed with PK in bulk before emulsification, and thermal activation triggers the release of mRNA for barcoding after emulsification.

**Figure 2.**
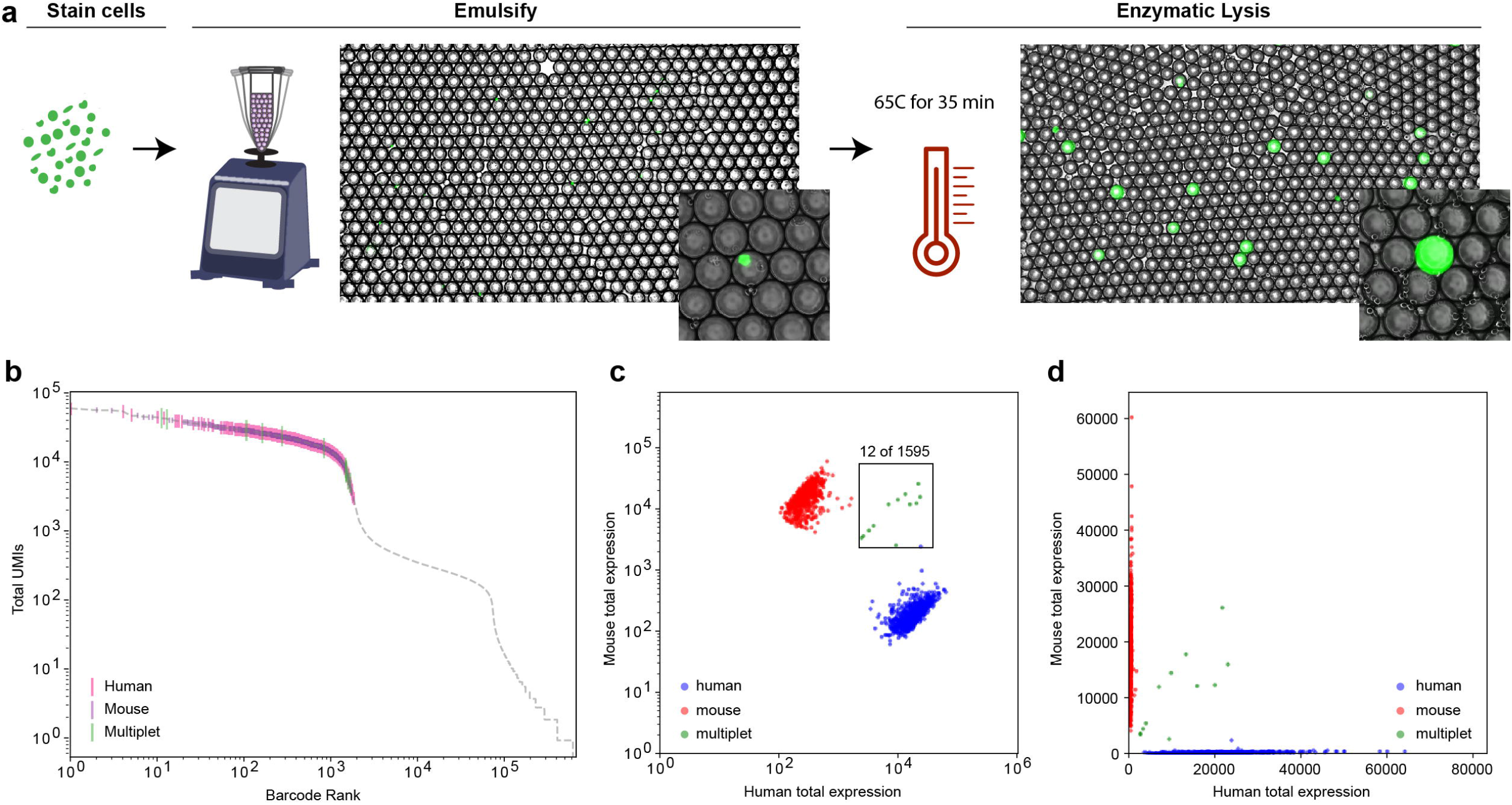
Heat-activated enzymatic lysis yields high purity single-cell transcriptomes. **(a)** Fluorescence microscopy (Brightfield, GFP) of calcein-stained cells emulsified with barcoded bead templates before and after heat-activated lysis. Inset: images show cell puncta (left) and release of calcein (right) after lysis. **(b-d)** Cell purity assessed with mouse-human mixing studies. **(b)** The distribution of total UMIs as a function of cell barcode rank. **(c-d)** Purity analysis of cell transcriptomes assessed using barnyard plots. Cells are colored by cell type: red points are mouse reads, blue points are human reads, and green points are mixed reads.

To ensure that temperature-activated lysis and bulk agitation do not pre-lyse cells and result in mRNA cross-contamination, we performed mouse-human cell line mixing studies. We synthesized barcoded polyacrylamide beads with polyT sequences, using split-pool ligation of four 6 bp randomers^30^. Beads contained ~10^8^ (96^4^) unique barcodes, providing ample sequence space to label a million cells. PIP-seq barcode rank plots for mixed mouse-human cell suspensions allowed cell identification by UMI abundance (**Fig. 2b**). The fraction of mouse reads in human transcriptomes was below 3%, and transcriptomes containing multiple cells were rare and consistent with Poisson encapsulation of two cells **(Fig. 2c,d)**. These results illustrate that PIP-seq yields high-purity single-cell RNA-seq data with minimal transcriptome mixing and low doublet formation.

### Accurate and scalable reconstruction of single-cell phenotypes in complex tissue

An important application of single-cell sequencing is atlasing cell types in heterogeneous tissue. To investigate the feasibility of atlasing studies, we applied PIP-seq to samples derived from healthy breast tissue. In tandem, we performed scRNA-seq on tissues from the same patients using a commercially available scRNA-seq technology (10x Genomics, Chromium v3). We integrated PIP-seq data across participants and recovered expected cell types by dimensionality reduction, including the two lineages of luminal epithelial cells (LEP1 and LEP2), myoepithelial cells, fibroblasts, vascular cells, and immune cells **(Fig. 3a, Extended data Fig. 3a,b)**^31^. To compare transcriptome capture between platforms, we downsampled the 10x Chromium and PIP-seq datasets to an equivalent number of cells and reads (2400 cells and 36,500 reads per cell). Chromium detected more unique genes (2298 *vs*. 1757, median) and transcripts (7491 *vs*. 3394) per cell, with similar percentages of reads assigned to mitochondrial transcripts (2.34% *vs*. 1.32%) **(Extended data Fig. 3c)**. To compare the transcriptome accuracy of PIP-seq, we downsampled each dataset to an equivalent number of unique molecular identifiers (UMIs) per cell (2400 cells, 1500 UMIs), integrated the data, performed dimensionality reduction, and identified clusters **(Fig. 3b,c)**. We compared marker genes and the correlation between gene expression profiles by cluster. Predicted marker genes were concordant between methods **(Fig. 3d)**, gene expression was highly correlated **(Fig. 3e, Extended data Fig. 4a)**, and breast tissue markers from previous reports were segregated identically within integrated clusters **(Extended data Fig. 4b).** Comparison of PIP-seq to publicly available data from 10x (v3, v2) and previously published scRNA-seq workflows demonstrated that PIP-seq produced high-quality transcriptomes across a range of sequencing depths **(Extended data Fig. 5)**. Next, we validated the scalability of PIP-seq, capturing and scRNA sequencing 138,146 breast tissue cells in a single tube reaction, as well as 65,000 peripheral blood mononuclear cells (PBMC) **(Extended data Fig. 6a-c)**. At high cell numbers, we identified a population of CD34 hematopoietic stem/progenitor cells in the PBMC sample, highlighting the importance of scalability in detecting rare cell types **(Extended data Fig 6b,c).** Last, we validated that PIP-seq is compatible with antibody-based cell hashing **(Extended data Fig 6d,e)**. Hashing can be used to further increase the number of cells and conditions processed. Thus, PIP-seq is an easy-to-use, accurate and scalable method to profile complex tissues.

**Figure 3.**
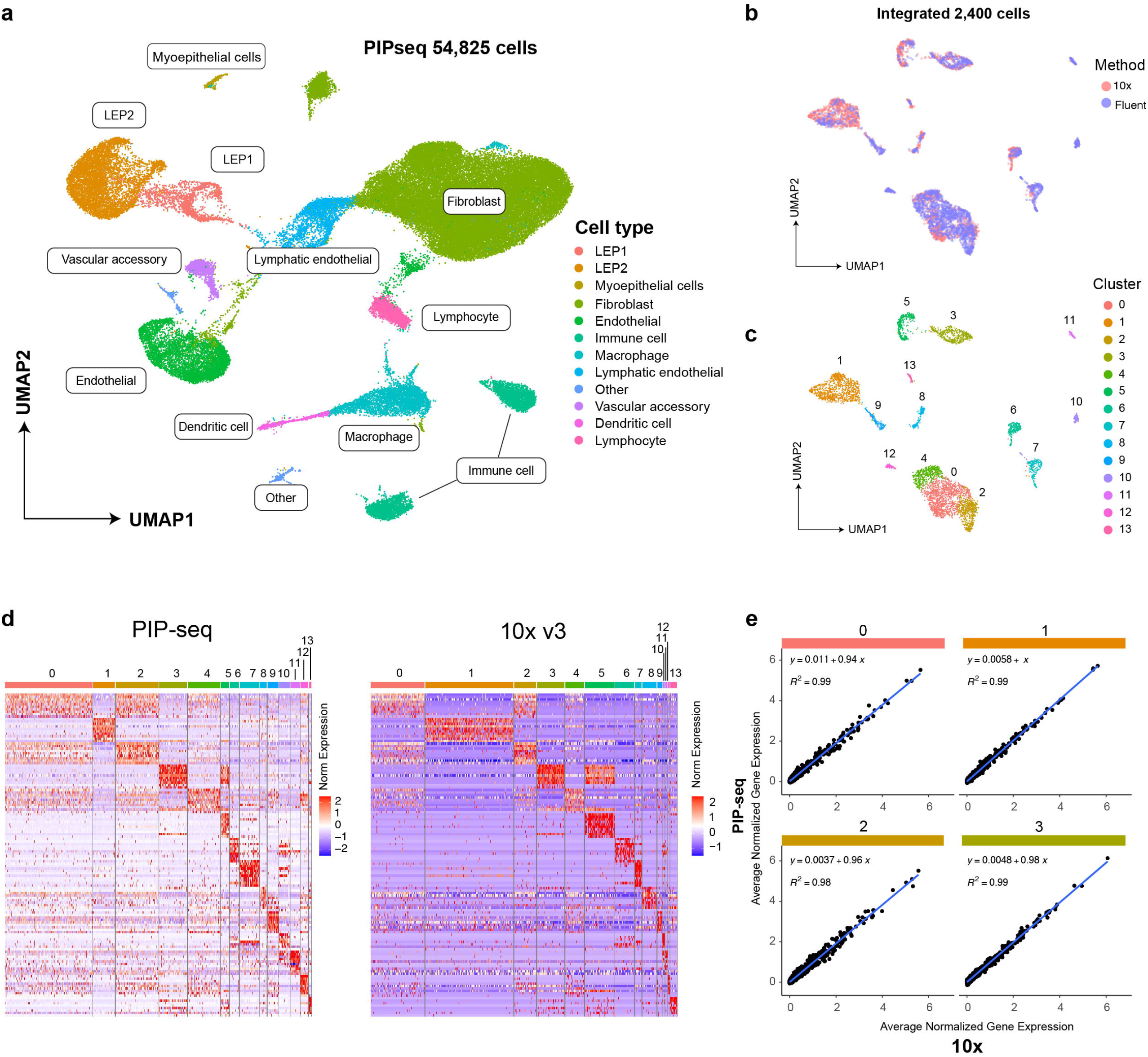
Accurate single-cell transcriptional profiling of healthy breast tissue using PIP-seq. **(a)** Clustering and identification of cell types from PIP-seq data (54,825 cells from 2 patients). **(b-e)** Comparison of PIP-seq to 10X Genomics data collected from the same tissue. **(b)** Integration of PIP-seq and 10x data. **(c-d)** Cell clustering and comparison of marker genes between platforms. **(d)** Heatmaps of marker gene expression show similar patterns in PIP-seq and 10X data. **(e)** Correlations in normalized gene expression, by cluster, between platforms (see also Extended data Fig. 4a).

### PIP-seq for single-cell pooled CRISPR screens

CRISPR perturbations combined with single-cell sequencing allow unbiased discovery of genotype-phenotype relationships^32–34^. Expanding this approach to genome-wide sgRNA libraries can elucidate gene function on an unprecedented scale. However, such studies require sequencing millions of cells to characterize all perturbations in libraries with tens or hundreds of thousands of individual sgRNA^19^. To demonstrate how the throughput of PIP-seq enables perturbation studies at scale, we profiled the transcriptional changes associated with a CRISPR interference allelic series CROP-seq library^35^. This library expressed sgRNA and a polyadenylated copy of the guide sequence from separate promoters. Guide RNAs were captured and barcoded with the cell’s polyadenylated mRNA, making this approach immediately compatible with PIP-seq. The library is designed to quantitatively titrate gene expression using sgRNAs with target site mismatches^35^, allowing us to compare measured gene expression to expected knockdown efficiency across each gene’s allelic series **(Fig. 4a).** We transduced K562 cells containing a stable dCas9-KRAB with the CRISPRi lentiviral library and performed PIP-seq to capture the transcriptional profiles and sgRNA identity of individual cells **(Fig. 4b-c)**. For cells with single gRNA assignments, previously reported knockdown efficiencies^35^ correlated with the normalized counts of targeted genes **(Fig. 4d)** and were most significant for highly expressed genes **(Extended data Fig. 7a,b).** In addition, the knockdown of genes produced known transcriptional changes. For example, gRNA targeting HSPA5 resulted in endoplasmic reticulum stress and increased unfolded protein response **(Fig. 4e).** These results validate the use of PIP-seq for CROP-seq experiments, paving the way for routine million-cell experiments that map genotype-phenotype relationships at genome-scale.

**Figure 4.**
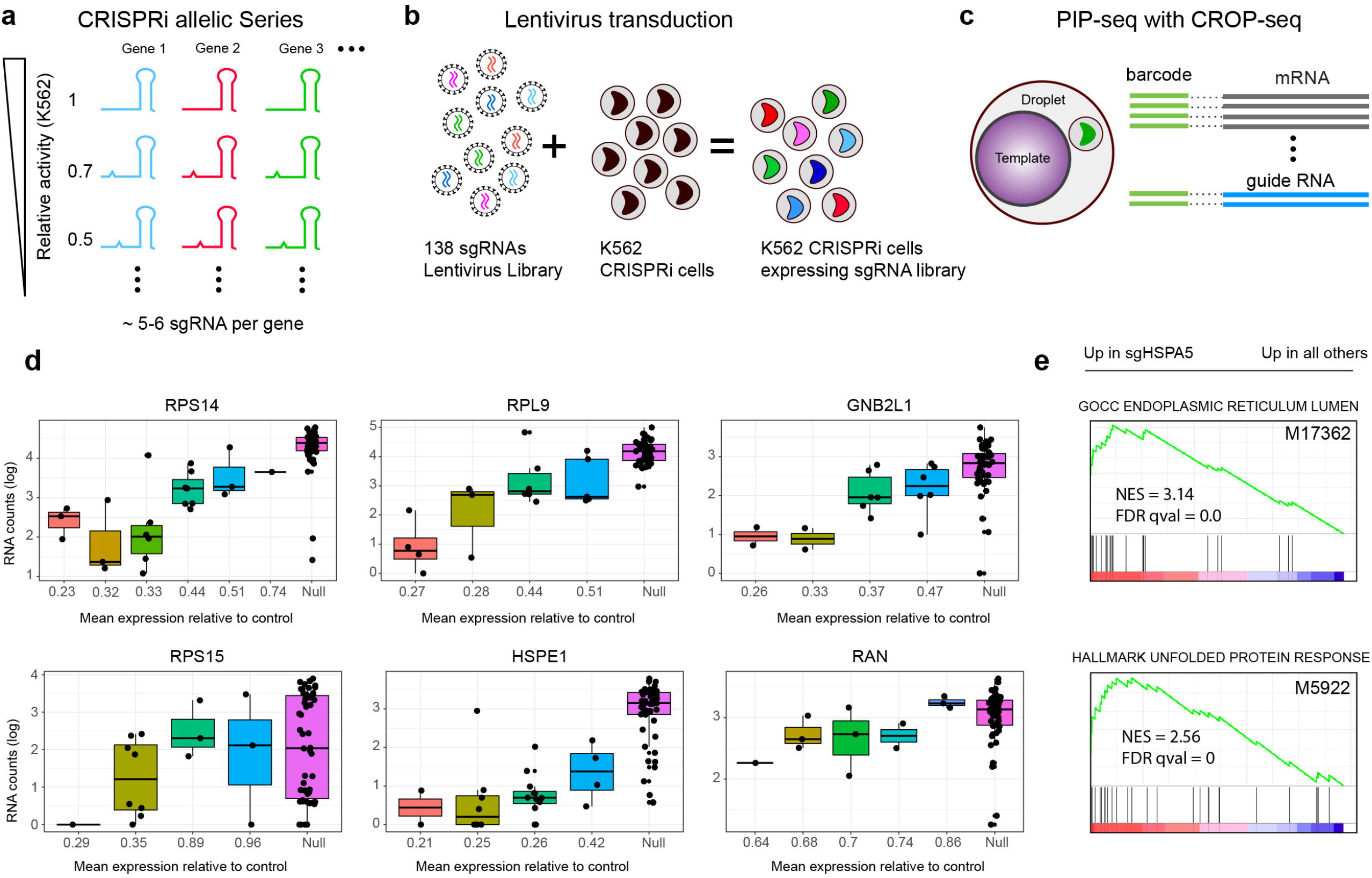
Transcriptome and guide RNA sequencing using PIP-seq. **(a)** Schematic of the CROP-seq sgRNA library designed with target mismatches to modulate the activity of essential genes. **(b)** Lentiviral transduction of the CRISPRi library in K562 cells. **(c)** Schematic of the capture and barcoding of polyadenylated mRNA and sgRNA using PIP-seq. RNA and sgRNA libraries are prepared separately and pooled for sequencing. **(d)** Quantification of gene expression of sgRNA within an allelic series. sgRNA is ordered from high to low predicted knockdown efficiency (Jost et. al, 2020). Non-targeting sgRNA are denoted as “Null.” **(e)** Pre-ranked gene set enrichment analysis (GSEA) of scRNA-seq data comparing sgHSPA5 transduced cells to non-sgHSPA5 transduced cells shows enrichment in genes related to endoplasmic reticulum stress and unfolded protein response.

### PIP-seq identifies unique single-cell transcriptomic signatures correlated with the relapse of mixed phenotype acute leukemia (MPAL)

Monitoring of cancer in response to therapy is an emerging application of single-cell sequencing that benefits from rapid sample processing at the point of collection, and the ability to delay cDNA synthesis and library preparation until multiple samples have been collected. We investigated the utility of PIP-seq for understanding cancer dynamics by first validating the single-cell transcriptional responses of two cancer cell lines (H1975, PC9) to Gefitinib, an epidermal growth factor receptor (EGFR) tyrosine kinase inhibitor. We treated H1975 and PC9 with DMSO (vehicle control) or 1 μM Gefitinib overnight and performed PIP-seq **(Fig. 5a)**. A transcriptional response in H1975, which is resistant to Gefitinib due to EGFR mutations L858R and T790M, was not observed, while Gefitinib sensitive PC9 cells showed a substantial shift in gene expression **(Fig. 5b).** Differential gene expression analysis revealed increased levels of *TACSTD2* (*TROP2*) in PC9 cells, consistent with its known modulation during lung adenocarcinoma tumor growth^36^ **(Fig 5c)**, and decreased expression of *CDK4*, which is known to enhance sensitivity to EGFR inhibitors^37^ (**Extended data Fig. 8)**. In addition, drug-resistant H1975 cells spiked into a background of sensitive cells (1:9 H1975:PC9) could be detected solely by their single-cell phenotypes, and at roughly the expected frequency (4.7%) **(Fig. 5d)**. Thus, PIP-seq recovered genes with reported roles in lung-cancer drug resistance and could identify resistance phenotypes within a background of drug-sensitive cells.

**Figure 5.**
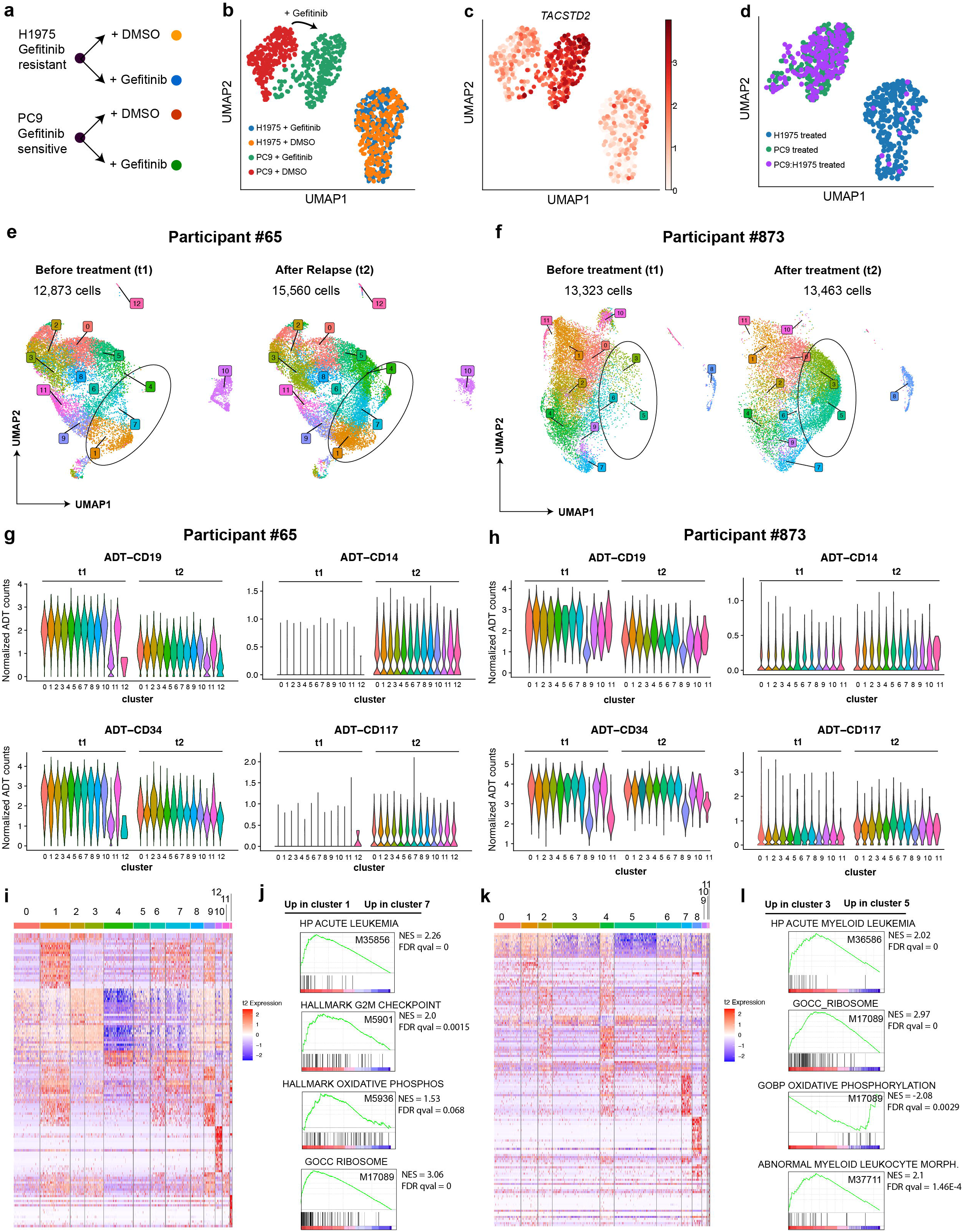
Molecular signatures of drug-resistant cancer phenotypes in cell lines and patient samples. **(a)** A two-by-two experimental study design using lung adenocarcinoma cell lines (H1975, PC9) treated with Gefitinib or DMSO. **(b)** Clustering of single-cell RNA-seq after drug treatment shows transcriptional perturbations in Gefitinib-sensitive PC9, but not Gefitinib-resistant H1975 cells. **(c)** Increased expression of *TACSTD2* in PC9 challenged with Gefitinib. **(d)** Identification of drug-resistant H1975 cells spiked into drug-sensitive PC9 cells based on Gefitinib-induced transcriptional perturbation. **(e-l)** PIP-seq RNA and barcoded antibody (CITE-seq) analysis of mixed phenotype acute leukemia (MPAL). **(e)** Clustering of single cells for patient 65 before (left panel) and after (right panel) chemotherapy. (**f)** Clustering of single cells for patient 873 before (left panel) and after (right panel) chemotherapy. **(g-h)** Antibody derived tag (ADT) abundance, by cluster, before (t1) and after (t2) chemotherapy. ADTs change as a function of chemotherapy but are consistent among clusters for both patients, with the exception of T cell subsets. **(i-l)** Analysis of transcriptional heterogeneity in MPAL samples **(i)** Heatmap of top differentially expressed marker genes by cluster after relapse in patient 63. **(j)** GSEA pre-ranked analysis comparing transcriptomic differences between clusters 1 and 7 in patient 65 using gene sets: HP Acute Leukemia (M35856), Hallmark G2M Checkpoint (M5901), Hallmark Oxidative Phosphorylation (M5936), and GOCC Ribosome (M17089) **(k)** Heatmap of top differentially expressed marker genes by cluster after relapse in patient 873. **(l)** GSEA pre-ranked analysis comparing transcriptomic differences between clusters 3 and 5 in patient 873 using gene sets HP Acute Myeloid Leukemia (M36586), GOCC Ribosome (M17089), GOBP Oxidative Phosphorylation (M17089), and Abnormal Myeloid Leukocyte Morphology (M37711).

Next, we applied PIP-seq to study mixed phenotype acute leukemia (MPAL), a high-risk disease characterized by multiple hematopoietic lineages^38,39^. Recurrence and changes in immunophenotype with chemotherapy are typically monitored using flow cytometry of surface markers during diagnosis, treatment, and relapse, but this provides limited insight into the drivers of relapse after drug treatment. Like other single-cell RNA-seq methods, PIP-seq can be multiplexed to simultaneously characterize single-cell gene expression and surface immunophenotype^40^. Using PIP-seq, we performed antibody-derived tag (ADT) sequencing (CITE-seq) on longitudinal samples collected from MPAL patients treated with chemotherapy. PIP-seq confirmed the diagnosis of these samples as B/myeloid MPAL and identified aberrant expression of immune and stem cell markers that matched with clinical immunophenotypes determined by flow cytometry **(Supplemental Table 2, Extended data Fig. 9)**. However, PIP-seq revealed an additional layer of complexity undetectable by traditional immunophenotyping. Dimensionality reduction identified new cell clusters that emerged after drug treatment **(Fig. 5e,f, Extended data Figs. 10, 11)**. These clusters had similar immunophenotypes **(Fig. 5g,h)**, but contained significant transcriptional heterogeneity **(Fig. 5i,k, Supplemental Tables 4,5)**. Cell populations up-regulating genes and pathways (oxidative phosphorylation, G2M checkpoint modulation, ribosome biogenesis) implicated in a variety of cancers including acute lymphoblastic leukemia^41–50^, but not previously linked to MPAL, were observed **(Fig. 5j,l)**. Taken together, our results highlight the value of single-cell methodologies for studying the heterogeneous response of cancer subpopulations to chemotherapy and the potential for the integration of simple and reliable scRNA-seq workflows into clinical research.

## Discussion

Genomics has progressed rapidly to high throughput, multi-modal, single-cell analysis^40,51–54^. Further improvements in data quality, the ability to measure additional cellular properties, and new computational approaches for understanding and integrating single-cell information^55–57^ will continue to refine our understanding of cell states. At the same time, there remains an unmet need for simplified workflows that scale in cell number and sample size, and that allow for breaks in processing after initial sample collection. PIP-seq is a microfluidics-free, single-cell RNA-sequencing method that produces high-quality data using a drastically simplified emulsification technique. Like other high-throughput single-cell approaches, PIP-seq is fundamentally a strategy to barcode mRNA from cells so that material can be pooled and sequenced. The core advantage of PIP-seq is the speed and simplicity of sample processing. Particle-templated emulsification forms monodispersed bead-containing emulsions in minutes with a standard laboratory vortexer, removing the need for instrumentation located in core facilities or hours of multichannel pipetting to perform split-pool indexing in plates. This expands access to single-cell technologies in several important ways. It reduces the need for sample transport, enabling immediate processing by technicians without prior training, and collection and banking of samples from remote locations, including field sites. It allows infectious samples that require special precautions to be processed at the point of collection or in the biosafety facilities where they are stored. More generally, rapid sample processing eliminates the need for fixatives and minimizes transcriptional perturbations and batch artifacts associated with processing many samples in series.

In addition to workflow simplicity, PIP-seq is intrinsically scalable, handling cell inputs over five orders of magnitude (10-10^6^), making it well suited for screening, genome-wide Perturb-seq experiments, and large cell-atlas studies. While methods based on combinatorial indexing scale efficiently to large cell numbers, PIP-seq has a substantially simpler workflow and is also compatible with high throughput processing of samples in plates, allowing many conditions and replicates to be run simultaneously. This has important implications for data quality and biological discovery in single-cell experiments since the detection of true positives and reduction in false positives in differential expression analysis is substantially improved by incorporating replicates and statistical methods that account for biological variability^58^. Increased flexibility in the number of samples that can be processed also enables previously difficult experimental designs, like dose-response curves, time-course studies, combinatorial perturbations, single-cell sequencing of organoids, and large drug screens. In addition, because PIP-seq can directly emulsify in plates, it integrates with robotic fluid handling and therefore comprises a drop-in solution for single-cell readouts in high-throughput experiments in academia or industry.

We confirmed the accuracy of PIP-seq as a single-cell genomics tool by profiling heterogeneous tissue and directly comparing our results to a commercial scRNA-seq platform (10x Genomics). PIP-seq cell type classification, marker identification, and gene expression levels were tightly matched with 10x data but detected fewer genes per cell. We attribute these differences to the extensive optimization that the commercial platform has undergone and suspect that, like other single-cell techniques^3–6,12,25,59^, further improvements to PIP-seq molecular biology will increase sensitivity. In addition, because PIP-seq emulsions are functionally equivalent to those made with microfluidics, our approach is immediately compatible with emerging advances, including improvements to the molecular biology of myriad multi-omic profiling methods developed for other droplet microfluidic barcoding systems^40,57,60^.

Finally, we demonstrated the utility of PIP-seq in processing clinical samples. In combination with barcoded antibodies, we profiled the relapse of mixed phenotype acute leukemia (MPAL) after chemotherapy. MPAL is a subtype of leukemia characterized by poor prognosis^61^, lineage ambiguity, lack of consensus regarding therapy, and significant intratumoral genetic and immunophenotypic heterogeneity^62,63^. The molecular mechanisms underlying treatment resistance in this complex disease remain undefined. Changes in gene expression have been linked to prognosis and treatment resistance in multiple cancers. However, tumor heterogeneity makes it unlikely that bulk sequencing methods would identify strong gene signatures associated with resistance in clinical samples. Using PIP-seq of longitudinal samples from two individuals with MPAL with disease progression after initial therapy, we identified transcriptional heterogeneity beyond that observed by immunophenotype and speculate that this heterogeneity may play a role in MPAL treatment resistance. We observed up-regulation of genes and pathways previously associated with acute lymphoblastic leukemia in several cell subsets that emerged after chemotherapy, as well modulation of ribosomal genes in both patients. Control of translation has been previously implicated in many cancers^41–46,64^, including leukemia, but has not yet been linked to MPAL progression and drug resistance, suggesting that new therapeutics targeting ribosomal biogenesis and/or protein translation may also have therapeutic potential in MPAL^65^. Our results motivate the use of single-cell technologies for understanding MPAL tumor heterogeneity and response to chemotherapy and suggest that the broad adoption of such technologies for monitoring cancer progression (and tailoring treatment) is within reach. In summary, single-cell RNA-sequencing provides unparalleled insight into cell heterogeneity but remains underutilized in many settings. PIP-seq addresses this with a simple, rapid, and scalable workflow that can be used by any lab containing standard molecular biology equipment.

## Methods

### Proteinase K triggered cellular lysis and mRNA capture

Mammalian cells were stained with Calcein AM (ThermoFisher #C3099) in 1 mL of PBS with 0.04% BSA according to manufacturer’s instructions. After 30 min of incubation at room temperature on a rotisserie incubator (Isotemp, Fisher Scientific), cell suspensions were quantified with a Luna-FL automated cell counter and diluted in 1X PBS with 0.04% BSA. 1500 calcein-stained cells in 5 μL of were added to 35 μL of barcoded hydrogel templates with 29 U/mL Proteinase K (NEB #P8107S) and 70 mM DTT (Sigma #D9779) and mixed for 10 pipet strokes. Care was taken to avoid generating bubbles when mixing cells with 280 μL of 0.5% ionic Krytox in HFE 7500 oil^66^ was added to the cell-bead mixture and vortexed at 3000 RPM for 15 seconds horizontally and then 2 minutes vertically with a custom vortexer (Fluent BioSciences, #FB0002776). Oil was removed from below the emulsion such that less than 100 μL remained. The PIP emulsion was subsampled on a C-Chip disposable hemacytometer (Fisher Scientific # DHCN015) before lysis with each subsample consisting of 3.5 μL of PIP emulsion per field of view. The C-chip was imaged in bright-field at 2X magnification. The remaining PIP emulsion was subjected to enzymatic lysis at 65°C for 35 min on a PCR thermocycler (Eppendorf Mastercycler Pro) with lid temperature set to 105°C. After lysis was complete, fluorescence images were captured using a Nikon 2000 microscope with 470 nm excitation (Thorlab M470L5).

### Synthesis of barcoded bead templates

Prototype barcode bead fabrication proceeded according to previous reports^30^. Briefly, a simple co-flow microfluidic device was used to combine acrylamide premix (6% w/v Acrylamide, 0.1% Bis-Acrylimide, 0.3% w/v ammonium persulfate, 0.1 □×□Tris-buffered saline–EDTA (TBSET: 10 mM Tris–HCL pH 8.0, 137 mM NaCl, 20 mM EDTA, 1.4 mM KCl, 0.1% v/v Triton-X100), 50 uM acrydited primer (/5Acryd/TTTTTTTAAGCAGTGGTATCAACGCAGAGTACGACTCCTCTTTCCCTACACGACGCTCTTCC) with oil (HFE-7500, 3 M Novec) containing 2% (w/v) surfactant (008-Fluoro-surfactant, Ran Technologies) and 0.4% v/v Tetramethylethylenediamine (TEMED). The emulsion was solidified at room temperature for 12 hours and beads were removed using 1H,1H,2H,2H-Perfluoro-1-octanol (Sigma Aldrich), washed thrice with Tris–EDTA-Tween buffer (TET: 10 mM Tris–HCl pH 8.0, 10 mM EDTA, 0.1% v/v Tween-20) followed by two washes with 30 mM NaCl, 10 mM Tris–HCl pH 8.0, 1 mM MgCl_2_, 0.1% Tween-20. The final bead size was 80 microns. Split-pool barcode assembly utilized the ligation assembly approach as described previously^30^. Beads were resuspended in T4 ligation buffer (NEB #B0202S), heated with a complementary oligonucleotide to 75°C for 2 min, and cooled to room temperature to anneal. 100 microliters of beads were distributed into each well of a 96-well plate containing a unique barcode with 1x T4 ligation buffer and 1.9 U/ul T4 DNA ligase (NEB #M0202M). Ligations were incubated at 25C for 1 hour and heat inactivated at 65C for 10 minutes. Well contents were combined and washed 5 times in 15mL TET. The process was repeated to add 4 barcodes and a UMI with polyT (NNNNNNNNNNNNTTTTTTTTTTTTTTTTTTTV). Quality control steps were identical to previous reports^30^. Bead manufacturing methods were transferred to Fluent BioSciences for scaled production, validation, and distribution. Commercially produced beads were used for several experiments, as noted.

### Varied format emulsification

PIP emulsification in varied formats was performed in 0.5 mL microcentrifuge tubes, 15 mL conical tubes and 50 mL conical tubes. Briefly, PIP particles were suspended in buffer with 29 U/mL Proteinase K Proteinase K (NEB #P8107S) and 70 mM DTT (Sigma #D9779) and pelleted through centrifugation. Barcoded hydrogel templates were then distributed at 35 μL, 0.5 mL and 8 mL volumes in 0.5 mL, 15 mL and 50 mL tubes, respectively. Fluorinated oil with surfactant (Fluent Biosciences # FB0001804) was added to each tube at 200 μL, 8 mL and 32 mL volumes, respectively. Emulsification was conducted on a Vortex Genie 2 with a custom adapter (Fluent #FBS-SCR-8VX) at maximum rpm for 1 min. After emulsification, the samples were allowed to settle for 30 seconds, and excess oil was removed via syringes using G22 blunt needles. The emulsion was subsampled, loaded on a C-Chip disposable hemacytometer (Fisher Scientific #DHCN015), and imaged under brightfield microscopy (DIAPHOT300, Nikon) at 2x and 4x magnification.

Emulsification in well plates was tested using two bead buffer conditions. First, to test emulsification in 96, 384, and 1536 well plates, PIP particles were suspended in 2% (v/v) Triton X-100 (Sigma, X100-5ML) in 10 mM TrisHCl (Teknova, T1075), then centrifuged at 6000 rcf, and the supernatant removed **(Fig. 1, Extended data Fig. 2c)**. Depending on the well plate working volume, 38 μL, 8 μL, or 3 μL of the centrifuged barcoded hydrogel templates were added to 96, 384, or 1532 well plates respectively. For 96 and 384 well plates 2 μL of sample was added to each well, for 1532 well plates, 1 μL was added to each well. PIP and sample volumes totaled 25% of the volume of each well. The plate was then sealed (Applied Biosystems, 4306311) and shaken for 5 min (IKA, 253614 and 3426400) to ensure complete mixing. Well plates were centrifuged at 200 rcf for 1 min before removing the seal. Then 80 μL, 20 μL, or 8 μL 2% (w/w) fluorosurfactant (Ran BioTechnologies, 008 Fluorosurfacant) in HFE oil (3M, Novec 7500) was added to each well in 96 (Applied Biosystems, N8010560), 384 (Applied Biosystems, A36931), or 1532 (Nunc, 253614) well plates respectively. The addition of oil represented 50% of the volume of each well for a total volume of 75% consisting of PIP, sample, and oil. After resealing, PIP emulsification was performed by vortexing for 30 seconds at 3200 rpm (Benchmark Scientific, BV1003). The emulsified plate was centrifuged at 200 rcf for 1 min before removing the seal and imaging droplets from individual wells on a fluorescence microscope (EVOS FL Auto).

Second, to test well plate emulsification with cells in 96 and 384 well plates, PIP particles were suspended in buffer with 29 U/mL Proteinase K (NEB #P8107S) and 70 mM DTT (Sigma #D9779) and pelleted through centrifugation. For 96-well plates (Eppendorf #0030129300), 25 μL of barcoded hydrogel templates were then distributed into each well with 4000 cells per well (2000 cells/μL x 2 μL). 150 μL of fluorinated oil with surfactant (Fluent Biosciences # FB0001804) was added to each well. Emulsification was conducted on a Vortex Genie 2 with flat-head adapter at 3000 rpm for 2 minutes. For 384-well plates (Corning #3347), 15 μL of barcoded hydrogel templates were then distributed into each well with 3000 cells per well (2000 cells/μL x 1.5 μL). 105 μL of fluorinated oil with surfactant (Fluent Biosciences # FB0001804) was added to each well. Emulsification was conducted on a Vortex Genie 2 with a flat-head adapter at 3000 rpm for 2 minutes **(Fig. 1, Extended data Fig. 2a,b)**.

### PIP-seq protocol

Unless otherwise noted, cells were centrifuged 300 x g for 5 min, washed twice in 1X PBS without calcium or magnesium (ThermoFisher #70011044) with 0.04% BSA, filtered with a 70 μm cell strainer, and resuspended in 1X PBS with 1% Pluronic F127 (Sigma #P2443). Pre-aliquoted barcoded hydrogel templates were thawed on ice. Volumes of barcoded hydrogel templates, cells, and oil varied based on the number of cells as noted in each experimental subsection below. A standard small-format run was as follows: 5 μL of 500 cells/μL was added to 35 μL of barcoded hydrogel templates with 29 U/mL Proteinase K and 70 mM DTT (Fluent BioSciences # FB0001876) and mixed for 10 strokes. Care was taken to avoid generating bubbles when mixing cells with barcoded hydrogel templates. 280 μL of oil (Fluent Biosciences # FB0001804) was added to the cell-bead mixture and vortexed (Vortex Genie 2, Scientific Industries) using a custom adapter (Fluent BioSciences, #FB0002100) at the maximum rpm for 15 seconds horizontally and 2 minutes vertically. 230 μL of excess oil was removed and the emulsion and enzymatic lysis was completed at 65°C for 35 minutes with a 4°C hold on a PCR thermocycler with lid temperature set to 105°C. The remaining oil was removed. The emulsion was broken as follows. Using a multichannel pipet, 180 μL of room temperature high salt buffer (250 mM Tris-HCl pH 8, 375 mM KCl, 15 mM MgCl_2_, 50 mM DTT) was added to the top of the emulsion followed by 40 μL of 100% 1H,1H,2H,2H-perfluoro-1-octanol (Sigma Aldrich, 370533). The samples were vortexed for 3 seconds, briefly centrifuged, and the bottom oil phase was removed. Barcoded hydrogel templates were transferred into a 1.5 ml Eppendorf tube and washed 3 times with 2x RT buffer (100 mM Tris-HCL pH 8.3, 150 mM KCl, 6 mM MgCl_2_, 20 mM DTT) with 1% Pluronic F68 (Gibco #24040032). After washing, the beads were pelleted, aqueous removed, and the remaining bead and buffer volume was 25 ul. To this mix, 25 μL of reverse transcription (RT) master mix. The master mix consisted of 4.8% PEG8000, 4% PM400, 2.5 μM template switch oligo (PIPS_TSO), 1 mM dNTPs (NEB), 1 U/μL RNase Inhibitor (NxGen, Lucigen), 1 U/μL reverse transcriptase (ThermoFisher, Maxima H-minus EP0751). RT master mix was added, mixed, and cDNA synthesis was completed for 30 minutes at 25°C, 90 minutes at 42°C, followed by 10 minutes at 85°C and a 4°C hold. Whole transcriptome amplification (WTA) was performed directly on RT product without purification by adding 50 μL of 1x KAPA Hifi master mix, 0.25 μM primer (PIPS_WTA_primer), and thermocycling 95°C for 3 min, then 16 cycles of (98°C 15 s, 67°C 20s, 68°C 4 min) followed by 72°C 5 min and 4°C hold. After WTA, barcoded hydrogel templates were removed using Corning Spin-X filter columns (1 min at 13,000 xg), and amplified cDNA was purified using 0.6x Ampure XP. Libraries were generated from WTA amplified material using the Nextera XT DNA Library Preparation Kit with a custom primer (PIPS_P5library) and standard Nextera P7 indexing primers (N70x). Libraries were pooled and sequenced using an Illumina NextSeq 2000 instrument with 15% PhiX. Oligonucleotides used in this study are found in **Supplemental Table 1**.

### Human-mouse mixing studies

Human HEK 293T cells (ATCC # CRL-3216) were grown in DMEM (ThermoFisher #11995073) supplemented with 10% FBS (ThermoFisher #A3840001) and 1% Penicillin-Streptomycin-Glutamine (ThermoFisher #10378016). Murine NIH/3T3 cells (ATCC # CRL-1658) were grown in DMEM (ThermoFisher #11995073) supplemented with 10% bovine calf serum (ATCC # 30-2030) and 1% Penicillin-Streptomycin-Glutamine. Cells were grown to a confluence of ~70% and treated with TrypLE Express with Phenol red (ThermoFisher #12605010) for 3 minutes, quenched with an equal volume of growth medium, and centrifuged at 5 min at 200 x g. The supernatant was removed, and the cells were resuspended in 1X DPBS without calcium or magnesium. Cells were diluted to their final concentration in 1X DPBS with 0.04% BSA and mixed evenly to create a 50:50 Human:Mouse mixture. Cell viability was evaluated using acridine orange/propidium iodide stain (Logos Bio # F23001) and quantified with a Luna-FL automated cell counter. Cells were processed using the PIP-seq protocol as described above.

### 72-hour hold experiments

5 μL of a 50:50 mixture of human HEK 293T cells and murine NIH/3T3 cells (800 cells/μL) was added to 35 μL of barcoded hydrogel templates Fluent BioSciences Part # FB0003067) with 29 U/mL Proteinase K and 70 mM DTT and mixed for 10 strokes. 280 μL of oil (Fluent Biosciences # FB0001804) was added to the cell-bead mixture, which was vortexed on a digital vortexer using a custom adapter (Fluent BioSciences #FB0002084) at 3000 rpm for 15 seconds horizontally and 2 minutes vertically. 230 μL of excess oil was removed and the emulsion was placed in a pre-heated digital dry bath at 66°C for 38 min and 4°C for 11 minutes. Control samples proceeded to emulsion breaking while 0°C hold samples were placed in an ice bucket in the 4°C refrigerator for 72 hours prior to breaking emulsions. Breaking, mRNA extraction, reverse transcription, WTA, and cDNA isolation, adapter ligation-based library preparation and Illumina sequencing were performed as previously described.

### Healthy breast tissue comparison to 10X

Fresh reduction mammoplasty tissue was processed as previously described^31,67^. Bulk mammary tissues were mechanically processed into a slurry and digested overnight with collagenase type 3 (200U/mL, Worthington Biochem CLS-3) and hyaluronidase (100U/mL, Sigma-Aldrich H3506) in medium containing charcoal:dextran stripped FBS (GeminiBio 100-119). The digested fragments were size filtered into a sub-40 micron fraction and an above 100-micron fraction and cryopreserved. For PIP-seq, cells were thawed and then resuspended in PBS + 0.04% BSA and passed through a 70-micron FlowMi cell strainer (Sigma # BAH136800070). For 10x Genomics data, the 100-micron fraction were thawed, then further digested with trypsin, followed by dispase (Stemcell Technologies #07913) and DNAseI (Stemcell Technologies #07469) digestion to achieve single-cell suspension. For PIP-seq, 20 μL of cells (1500 cells/μL in PBS + 0.04% BSA) was added to 200 μL of barcoded hydrogel templates (Fluent BioSciences, #FB0002617) and mixed for 10 strokes. 1000 μL of oil (Fluent Biosciences # FB0001804) was added to the cell-bead mixture and it was vortexed on a digital vortexer using a custom adapter (Fluent BioSciences, #FB0002100) at 3000 rpm for 15 seconds horizontally and 2 minutes vertically. 800 μL of excess oil was removed and the emulsion was placed on a pre-heated digital dry bath at 66°C for 38 min and 4°C for 11 minutes. Breaking, mRNA extraction, reverse transcription, WTA, and cDNA isolation was performed under standard conditions. Adapter ligation-based library preparation was performed according to manufacturer’s instructions (Watchmaker Genomics, #7K0019-024). Samples were sequenced on the Illumina NextSeq 2000, with four patient samples pooled per P3 cartridge and sequenced at a read depth of approximately 36,500 reads/cell. For 10X Genomics, cells from each patient were labeled with MULTIseq barcodes^13^, then pooled and stained with DAPI to be sorted for DAPI-live cells. Single-cell libraries were prepared according to the 10X Genomics Single Cell V3 protocol (v3.1 Rev D) with the standard MULTIseq sample multiplexing protocol. The libraries were sequenced on a NovaSeq S4 lane at a read depth of about 70,000 reads/cell. To compare platforms, we downsampled PIP-seq and 10x data, which had different numbers of cells and sequencing depth per cell. The PIP-seq data had 54,825 cells, sequenced at approximately 36,500 reads per cell, while the 10x data had 2420 cells sequenced at approximately 70k reads per cell. Data were downsampled to 2400 cells and 36,500 reads in R (downsampleReads, DropletUtils). For correlation and marker gene comparisons, data were downsampled to 2400 cells and 1500 UMIs in R (SampleUMI, Seurat v4.1.0). Markers used for breast tissue cluster cell type calling are found in **Supplemental Table 2**.

### Single-tube large format breast tissue study

PIP-seq was performed as previously described, except that cells were counted and diluted with PBS + 0.04% BSA to a concentration of 10,000 cells/μL. 40 μL of cell suspension was input into 800 μL of barcoded hydrogel templates (Fluent BioSciences Part # FB0003067). 4000 μL of oil (Fluent Biosciences # FB0001804) was added to the cell-bead mixture and vortexed on a digital vortexer using a custom adapter (Fluent BioSciences # FB0002659) at 3000 rpm for 15 seconds horizontally and 2 minutes vertically. Excess oil was removed using a 3 mL syringe with a G22 blunt bottom syringe needle. Lysis proceeded using 3300 μL of a lysis emulsion (Fluent BioSciences #FB0003039) added to the cell-bead emulsion. The mixture was placed in a pre-heated digital dry bath at 37°C for 45 min and 4°C for 10 minutes. Breaking, mRNA extraction, reverse transcription, WTA, and cDNA isolation were performed under the same conditions as described previously. Adapter ligation-based library preparation was performed according to manufacturer’s instructions (Watchmaker Genomics, #7K0019-024). 80 ng of cDNA was used to prepare four replicate library preparations which were pooled and sequenced on two Illumina NextSeq 2000 P3 cartridges at a read depth of 13,025 reads/cell, after concatenation.

### CROP-seq

K562 CRISPRi cells were cultured in RPMI-1640 (Gibco #11875093) with 10% FBS (Thermo Fisher Scientific, #10438026) and 1% penicillin/streptomycin (Thermo Fisher Scientific, #15140148) in an incubator at 37°C with 5% CO2. K562 CRISPRi cells were transduced with a lentivirus library containing 138 sgRNA^35^ at a multiplicity of infection of 0.1. Lentivirus-infected cells (BFP+) were sorted to high purity using a BD FACS Aria III (100 μM nozzle) and processed according to the PIP-seq scRNA-seq workflow. 3 μL of cells (333 cells/μL) was added to 28 μL of barcoded hydrogel templates with 29 U/mL Proteinase K and 70 mM DTT and mixed for 10 strokes. 150 μL of 0.5% ionic Krytox in HFE 7500 oil was added to the cell-bead mixture and it was vortexed at 3000 RPM for 1 minute on a Vortex Genie 2 with a custom tube adapter. cDNA was processed according to the standard PIP-seq protocol to obtain sequence ready libraries containing transcriptome information. To recover sgRNA sequences, we implemented an additional amplification step. We amplified 1 ng cDNA in a 50 μL reaction using of primers P5-PE1 (0.5uM) and Weissman_U6 (0.25uM) **(Supplementary Table 1)** with 1x Kappa HIFI. Reactions were thermocycled at 95°C for 3 min followed by 10 cycles of [95°C for 20s, 70°C for 30s [-0.2°C per cycle], 72°C for 20s], followed by 8 cycles of [95°C for 20s, 68°C for 30s, 72°C for 20s], followed by 72°C for 4 minutes, and 4°C forever. Library PCR product enriched in sgRNA sequences was purified with a double sided 0.5x/0.8x Ampure XP bead cleanup and the size was determined (Agilent Tapestation).

Transcriptome and sgRNA libraries were pooled at 20:1 before sequencing. Reads were first processed to extract sgRNA sequences. The bioinformatics pipeline was run using a custom index built from the full human transcriptome (GENCODE v32) and guide RNA sequences (salmon v1.2.0.). This approach led to the recovery of >14,000 unique gRNA counts across all cell-associated barcodes. Cells were assigned to gRNA groups using a previously reported approach^32^. Briefly, cells were classified as uniquely expressing a single gRNA species if the guide’s expression was at least 10-fold higher than the sum of all other gRNAs. Similarly, cells were classified as containing multiple gRNAs in cases where the difference was smaller than one. For the 581 single cells sequenced, 2 did not have any gRNA, 441 contained a single gRNA, and 138 contained multiple gRNA. Cell barcodes were processed using Seurat v4.1.0. All gRNAs in the list of features were excluded from the identification of variable transcripts (feature selection) and in subsequent stages of dimensionality reduction and clustering. To understand the relationship between gRNAs and mRNA expression, gRNAs were ranked according to their expected level of knockdown, reported previously^35^, and a generalized additive model was used to assess groupwise trends for each set of gRNAs.

### Lung adenocarcinoma cell line experiments

PC9 was obtained from the RIKEN Bio Resource Center (#RCB4455). H1975 was obtained from the American Type Culture Collection (#CRL-5908). Cells were cultured in RPMI-1640 (Gibco #11875093) with 10% fetal bovine serum (FBS), penicillin, and streptomycin in an incubator at 37°C with 5% CO2. 1 μM Gefitinib (Frontier Scientific, 501411677) or DMSO was added to culture flasks 24 hours before cells were harvested for processing. PC9 and H1975 were both treated with Gefitinib and DMSO. To perform the cell mixing study, Gefitinib treated H1975 and Gefitinib treated PC9 were mixed at a ratio of 1:9 H1975:PC9. 5 μL of cells (400 cells/μL) were added to 28 μL of barcoded hydrogel templates with 22.8 U/mL Proteinase K and 28 mM DTT and mixed for 10 pipet strokes. 150 μL of 0.5% ionic Krytox in HFE 7500 oil^66^ was added to the cell-bead mixture and it was vortexed at 3000 RPM for 1 minute on a Vortex Genie 2 with a custom tube adapter. Triplicate tubes of 400 cells were processed per treatment condition. Data was analyzed using Seurat v4.1.0.

### Healthy PBMCs

Cryopreserved PBMCs were obtained from a commercial provider (AllCells, Lot #3052467). Cells were thawed and prepared for PIP-seq as previously described in the MPAL study, except that the final cell dilution was made in 1X PBS + 0.04% BSA. *For the high cell count PBMC study*. PIP-seq was performed as previously described in the high cell number breast tissue study except that cells were counted and diluted with PBS + 0.04% BSA to a concentration of 4300 cells/μL and 44 μL of cell suspension was input into 800 μL of barcoded hydrogel templates (Fluent BioSciences Part # FB0003067). Cryopreserved PBMCs used for cell hashing were obtained from a commercial provider (AllCells, and Lot #3082436) and prepared for PIP-seq as described previously. *For the cell hashing study*. Cell staining and PIP-seq were performed according to PIPseq Single Cell Epitope Sequencing User Guide (FB0002079). Briefly, one million PBMCs were resuspended in 47.5 μL of Cell Staining Buffer (BioLegend #420201) and 2.5 μL TruStain FcX block (BioLegend #422301) was added prior to mixing and incubation for 10 min on ice. Next, 1 μg of TotalSeqA antibody was diluted in Cell Staining Buffer and 50 μL of this antibody dilution was added to the blocked cells prior to incubation on ice for 30 min. Stained cells were washed in Cell Staining Buffer three times and resuspended in 1X PBS+0.04% BSA at 2000 cells/μL. For the PIP-seq, 20 μL of this cell resuspension was added to 200 μL of barcoded hydrogel templates (Fluent BioSciences #FB0002617) and processed through PIP-seq.

### Mixed phenotype acute leukemia (MPAL)

Patients whose samples were used in this study were treated at the University of California San Francisco. Samples were collected in accordance with the Declaration of Helsinki under institutional review board-approved tissue banking protocols, and written informed consent was obtained from all patients. Sample clinical characteristics are found in **Supplemental Table 3**. Cryopreserved peripheral bone marrow mononuclear cell samples (PBMCs) were thawed by hand until approximately 85% of ice remained. Using a 5 mL serological pipette, 1 mL of 4°C defrosting media (DMEM with 20% FBS and 2 mM EDTA) was added dropwise to each sample and then, without disturbing the remaining ice pellet, the sample was carefully transferred dropwise to a pre-prepared 40 mL aliquot of 4°C defrosting media. This was repeated until the contents of the entire cryovial were transferred into the 50 mL conical of defrosting media. The sample was inverted 4-5 times and centrifuged at 114 x g for 15 min at 4°C with no brake. The supernatant was aspirated and 10 mL of room temperature RPMI 1640 with 1% Penicillin-Streptomycin-Glutamine was used to gently resuspend the cells. Cell clumps were manually removed, and if necessary cells were filtered through a 70 micron cell strainer into a fresh 50 mL conical. The sample was inverted 2-3 times and centrifuged at 114 x g for 10 minutes with low brake at room temperature. The supernatant was aspirated and cells were resuspended in an appropriate volume of 1X PBS + 5% FBS. Cells were quantified with AO/PI and viability evaluated on the Luna-FL. 1-2 million cells were aliquoted into a new 15mL conical and then centrifuged at 350 x g for 4 min at 4°C and then the supernatant was aspirated and the tube was placed on ice. 45 μL of cold Cell Staining Buffer (BioLegend, 420201) was added per million cells and resuspended gently. 5 μL Trustain FcX block (BioLegend, 422301) was added per million cells and gently mixed 10 times with a wide-bore pipette tip. Cells were blocked on ice for 15 minutes. A custom pool of 19 TotalSeq A antibodies was obtained from BioLegend. Immediately prior to use, antibodies were mixed and centrifuged at 10,000 x g for 4 min at 4°C. 4.6 μL of 0.5 μg/μL antibody pool was added per million of the blocked cells and gently mixed 10 times with a wide-bore pipette tip. The samples were incubated on ice for 60 minutes. Next, 3.5 mL of cold Cell Staining Buffer was added, gently mixed with a wide-bore pipette tip, and slowly inverted twice to mix. Cells were centrifuged at 350 x g for 4 min at 4°C and then the supernatant was removed. The addition of cold Cell Staining buffer was repeated twice for a total of 3 washes. After the final supernatant aspiration, stained cells were resuspended in 1X PBS with 0.04% BSA and mixed 5-10 times until cells were completely suspended without visible clumps. Cell concentration was determined with AO/PI and viability was evaluated on the Luna-FL. Final dilutions were made in 1X PBS with 0.04% BSA. 20 μL of cells was added to 200 μL of barcoded hydrogel templates (1000 cells/μL) and processed according to PIPseq Single Cell Epitope Sequencing User Guide (FB0002079). Marker genes identified for patients 65 and 873 are found in **Supplemental Tables 4 and 5**, respectively.

### PIP-seq bioinformatic analysis

Analysis of sequencing data was performed using custom scripts to generate gene expression matrices starting from processed FASTQ sequences. The pipeline comprises 4 basic steps: barcode identification and error correction, mapping to reference sequences, cell calling, and gene expression matrix generation. Briefly, after demultiplexing the sequencing data each read in the FASTQ is matched against a “whitelist” of known barcodes. Reads were matched with a hamming distance tolerance of 1, meaning that the barcode portion of a read can differ from a whitelist entry by one base and still be matched to that barcode. Reads that did not match any barcode in the whitelist were discarded from further analysis. Matching reads were output to a new, intermediate FASTQ file that was then used for mapping against an appropriate transcriptome reference. Reference transcriptomes matching the species of each sample were prepared using the Salmon *index* function with the default k-mer size of 31^68^. GENCODE references were used to build the transcriptome indexes including GRCh38.p13 for human, GRCm38.p6 for mouse, and the combination thereof for HEK/3T3 cell mixture studies. Following barcoding, Salmon *alevin* v1.2.0^69^ was used to map reads to the full transcriptome. The intermediate FASTQ generated during barcoding were provided as input into *alevin* along with a list of all whitelisted barcodes contained in raw reads. After mapping, data were output as UMI counts matrices (sparse matrix, gene list, barcode list) with dimensions of *all barcodes x all genes in index*. An in-house python implementation of emptyDrops^70^, a standard scRNA-seq method to separate putative cells from background, was then applied. A custom threshold for each experiment was set, beneath which no true cell barcodes were expected to fall. As with emptyDrops, an estimated ambient profile across all barcodes beneath that threshold was created. A p-value was computed by comparing the gene expression profile for each barcode above the threshold against the ambient profile. Barcodes with a statistically significant difference (Benjamini-Hochberg adjusted-p<0.001) from the ambient background profile were categorized as cell-containing barcodes. The *alevin* output matrices were then subset to only include called cell barcodes. Gene expression matrices were normalized prior to performing unsupervised clustering and UMAP dimensionality reduction. First, gene expression counts for each cell were divided by the total counts for that cell and multiplied by a scaling factor of 10,000. Finally, the data were transformed to natural-log scale using log1p(). The Seurat package (v4.1.0) was used to perform downstream clustering, marker gene determination, and visualization in R. Seurat’s FindClusters() and RunUMAP() commands were used with default settings.

For saturation curve comparisons, PIPseq and 10x samples were downsampled to matching depths of 5K-80K reads per called cell. Downsampling was performed using seqtk for PIPseq samples and using DropletUtils read10xMolInfo() function with a molecule_info.h5 file directly downloaded from the 10x website. Inflection point-based cell calling was used to standardize cell calls across platforms. Median transcripts/cell and genes/cell were calculated from the cell fraction of the resulting count matrices. For violin plot comparisons, samples were prepared to match the same processing configuration used by Ding et al (2020)^28^. First, samples were downsampled to 53K reads per called cell and trimmed to 50bp for read 2 prior to processing, sampling in the same manner described above. Each violin plot represents the cell fraction from a single replicate of an HEK/3T3 cell mixture, with human and mouse split out into separate plots.

Analysis of PBMC data for the high cell count study was performed using custom scripts as described above, until the completion of mapping. Cell calling, clustering, and differential expression were performed using PIPseeker v1.0.0 (Fluent Biosciences) in *reanalyze* mode using -force-cells 65000. Top differentially expressed genes from the PIPseeker graph-based clustering result were used to determine cell types by comparing to a reference gene list (Supplementary Table 7). Log-normalized expression for key genes (e.g. CD34) were overlaid on the UMAP projection to highlight markers associated with specific cell types (color bar in log10 scale). Analysis of PBMC data for the cell hashing study was performed using PIPseeker v1.0.0 in *count* mode using STAR (v2.7.10a) and the PIPseeker human reference (https://www.fluentbio.com/products/pipseeker-for-data-analysis/). ADT analysis was performed by performing barcode error correction with PIPseeker v1.0.0 (*count* mode) and custom scripts to trim R2 to the first 16bp. Error-corrected and trimmed FASTQs were input to CITE-seq Count (v1.4.3) using the following settings: -t (hashtag whitelist) -cbf 1 -cbl 16 -umif 17 -umil 28 --cells (# called cells from RNA cell calling). The hashtag whitelist contained two TotalSeq™-A anti-human Antibody hashes (A0253 - TTCCGCCTCTCTTTG, A0255 - AAGTATCGTTTCGCA). The filtered matrix output by PIPseeker for the RNA data was merged with the umi-count matrix from CITE-seq Count on cell barcode to create a merged matrix. The hashing data was demultiplexed in Seurat using HTODemux (positive.quantile=0.99). Downstream was performed in Seurat using SCTransform() along with RunPCA(), FindNeighbors(dims=1:15) and RunUMAP(dims=1:15). Cell type annotation was performed with singleR (v1.4.1) and used an annotated 10x Genomics v1 chemistry dataset as a reference. Cells were classified by their max hash identity and projected in the RNA-based UMAP space. The HTO data was subjected to clustering in Seurat using the HTOHeatmap() function to visualize singlets, doublets, and unclassified cells.

For 72-hour hold experiments, analysis was performed using custom scripts as previously described. Samples were normalized to the same depth (45,000 reads/cell). Cell types were then annotated as human (HEK293T) or mouse (NIH-3T3) using a purity threshold of >85% single-species content per barcode. Barcodes from each species were subset and transcript counts were summed for each gene to generate two pseudobulk counts tables per sample. Samples were aggregated separately for each species and then were analyzed with DESeq2. A contrast of 0 vs. 72 hours was performed for each species, while controlling for batch effects associated with different users. For the correlation analysis, pseudobulk counts derived above were normalized to transcripts per million and transformed to log1p scale. Pearson correlations (R) and slopes (m) were calculated by fitting a linear model to the data. Data were then plotted in R with ggplot2 v3.3.5 and were aggregated into a grid using GGally v2.1.2. Additionally, the distribution of cells in UMAP space at 0 and 72 hours post-lysis was examined. After processing data in Seurat as described, harmony batch-correction was used to integrate datasets.

## Figures

**Extended data Figure 1.**
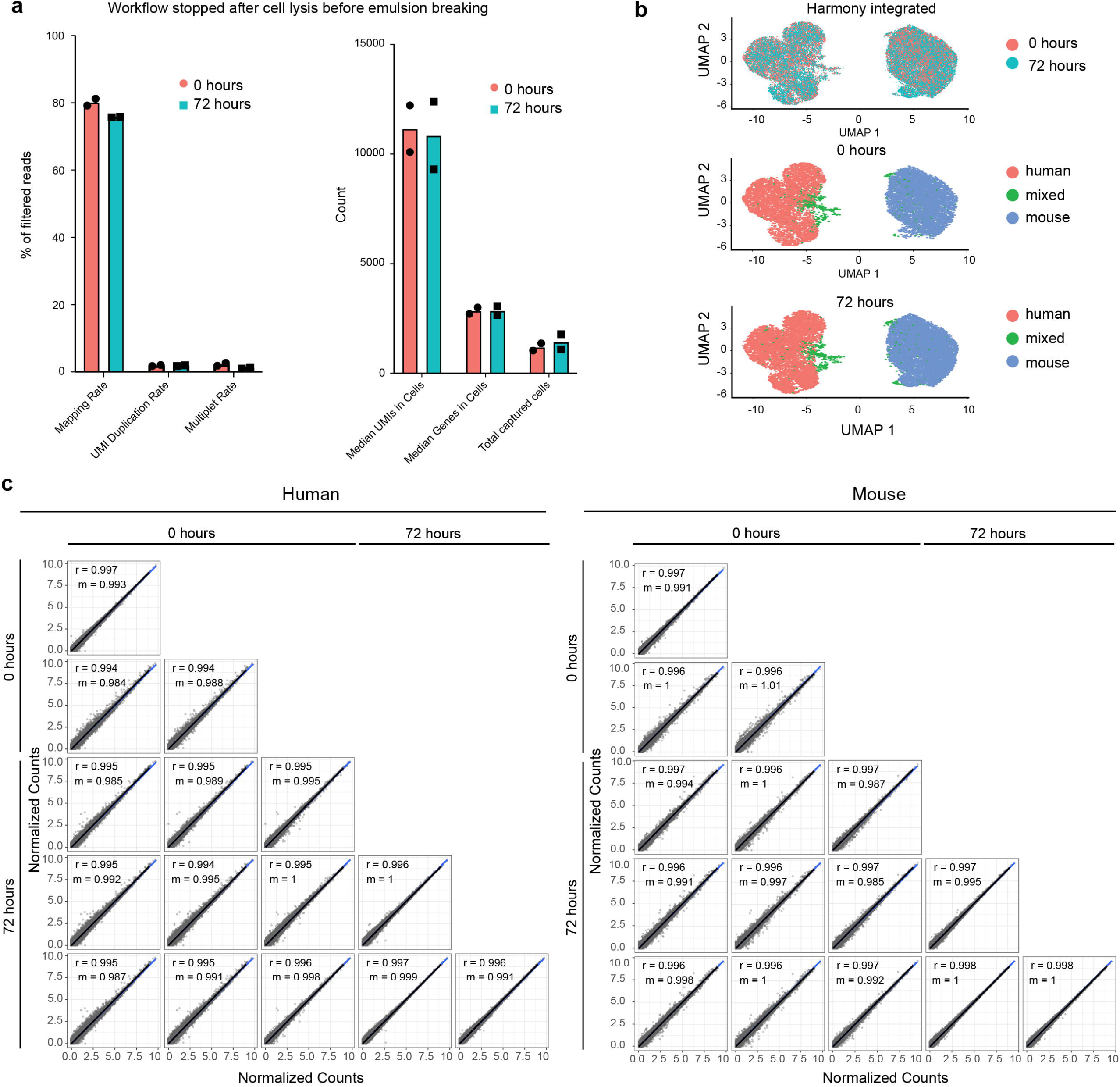
Flexible sample processing with PIP-seq. **(a)** Storage of droplets after emulsification for 72 hours at 0°C did not change quality metrics. **(b)** Data integration between time points. **(c)** Correlations in normalized gene expression, by sample, between time points for mouse and human cells.

**Extended data Figure 2.**
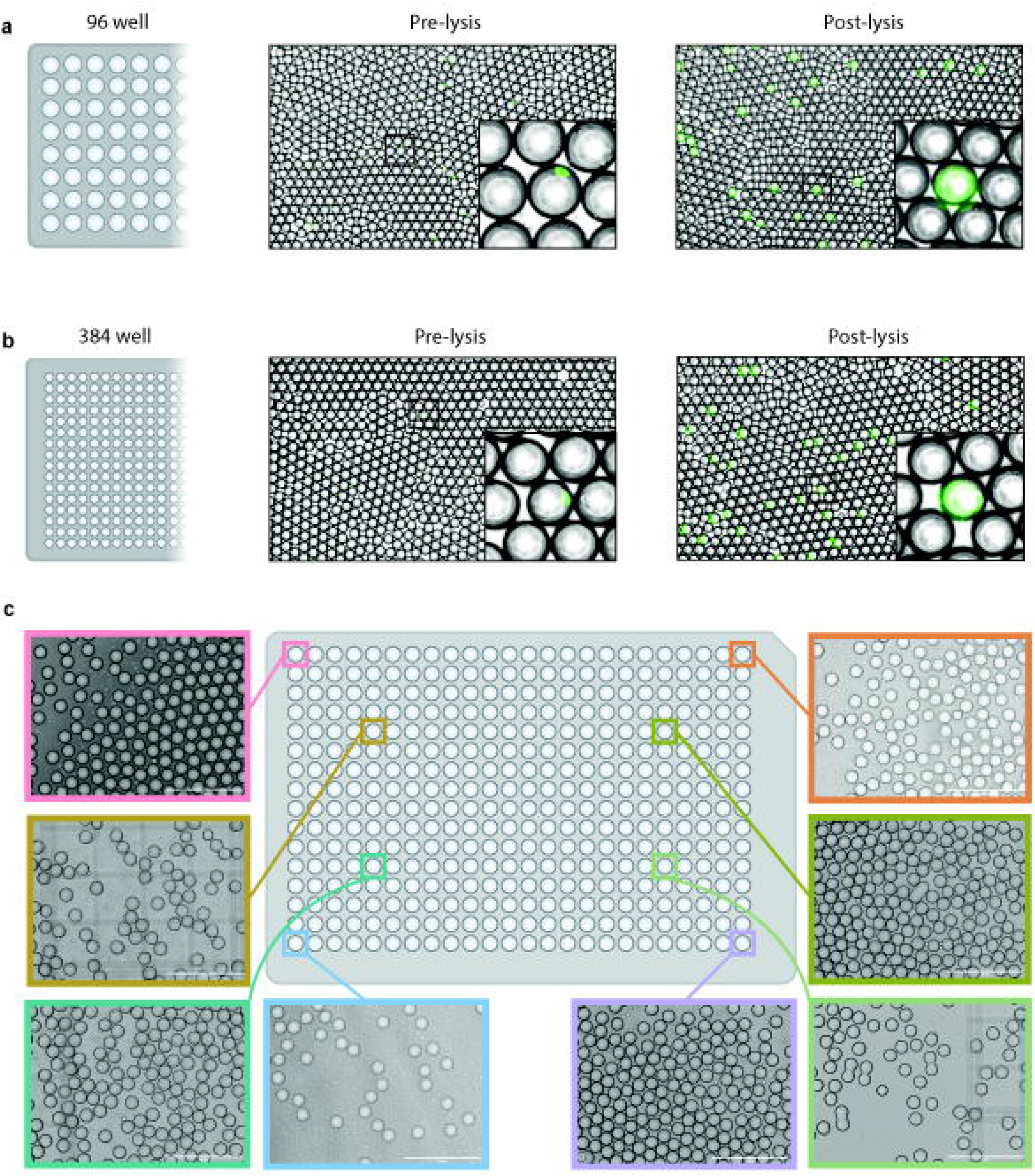
**(a-c)** Microscope images of droplets and cells in plate emulsification experiments. **(a,b)** Barcode bead templates, stained cells (puncta), and lysis reagents are combined with oil and vortexed in **(a)** 96-well and **(b)** 384-well plates to generate monodispersed droplets. Heat activation of Proteinase K results in lysis and release of calcein dye, and full-drop fluorescence. **(c)** Microscope images of droplets from random wells of a 384-well plate emulsification experiment.

**Extended data Figure 3.**
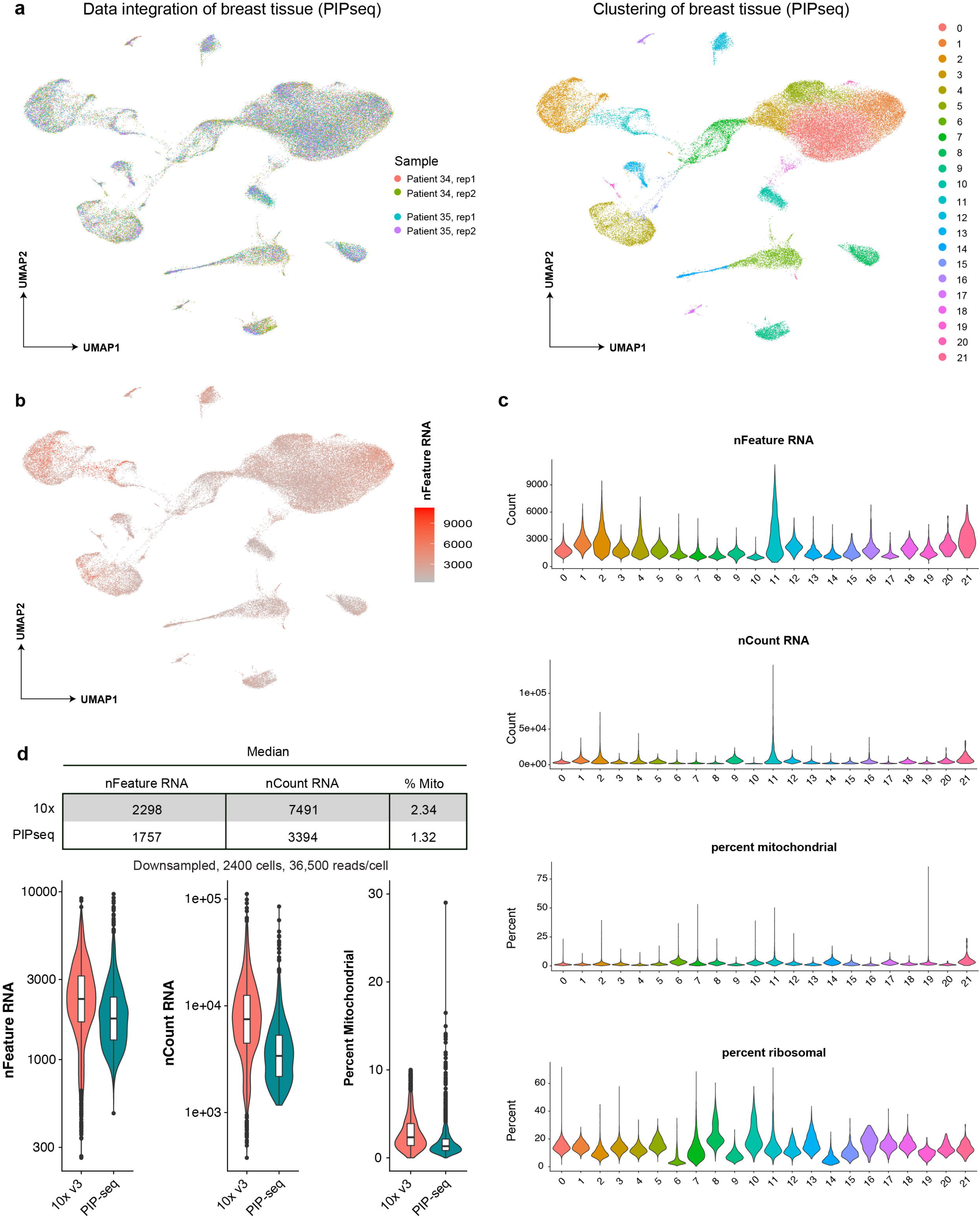
Quality control analysis of PIP-seq using healthy breast tissue. **(a)** Integration and clustering of 54,825 cells from 2 patients with 2 replicates per patient. **(b)** Coloring of UMAP by the number genes (nFeature RNA) for each cell. **(c)** The number of unique genes (nFeature RNA), transcripts (nCount RNA), percent mitochondrial reads, and percent ribosomal reads as a function of cluster. **(d)** Comparison between 10X Genomics’ and PIP-seq data after downsampling 2400 cells to 36,500 reads per cell.

**Extended data Figure 4.**
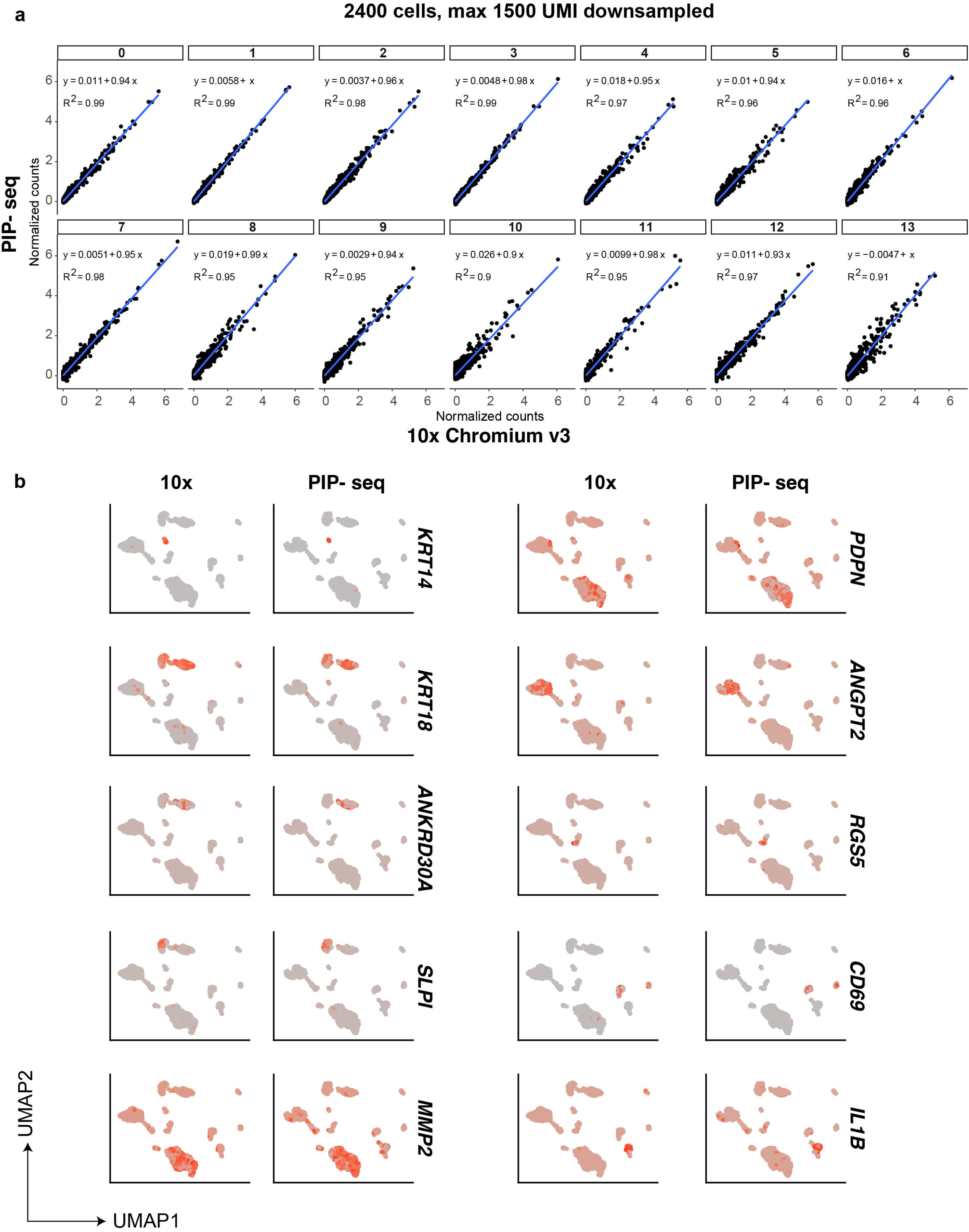
Comparison, after down-sampling (2400 cells 1500 UMIs) of the larger breast tissue PIP-seq dataset to 10x Genomics data generated from identical samples. **(a)** Correlations in normalized gene expression, by cluster, between platforms. **(b)** Expression of marker genes overlayed on clusters is consistent between platforms.

**Extended data Figure 5.**
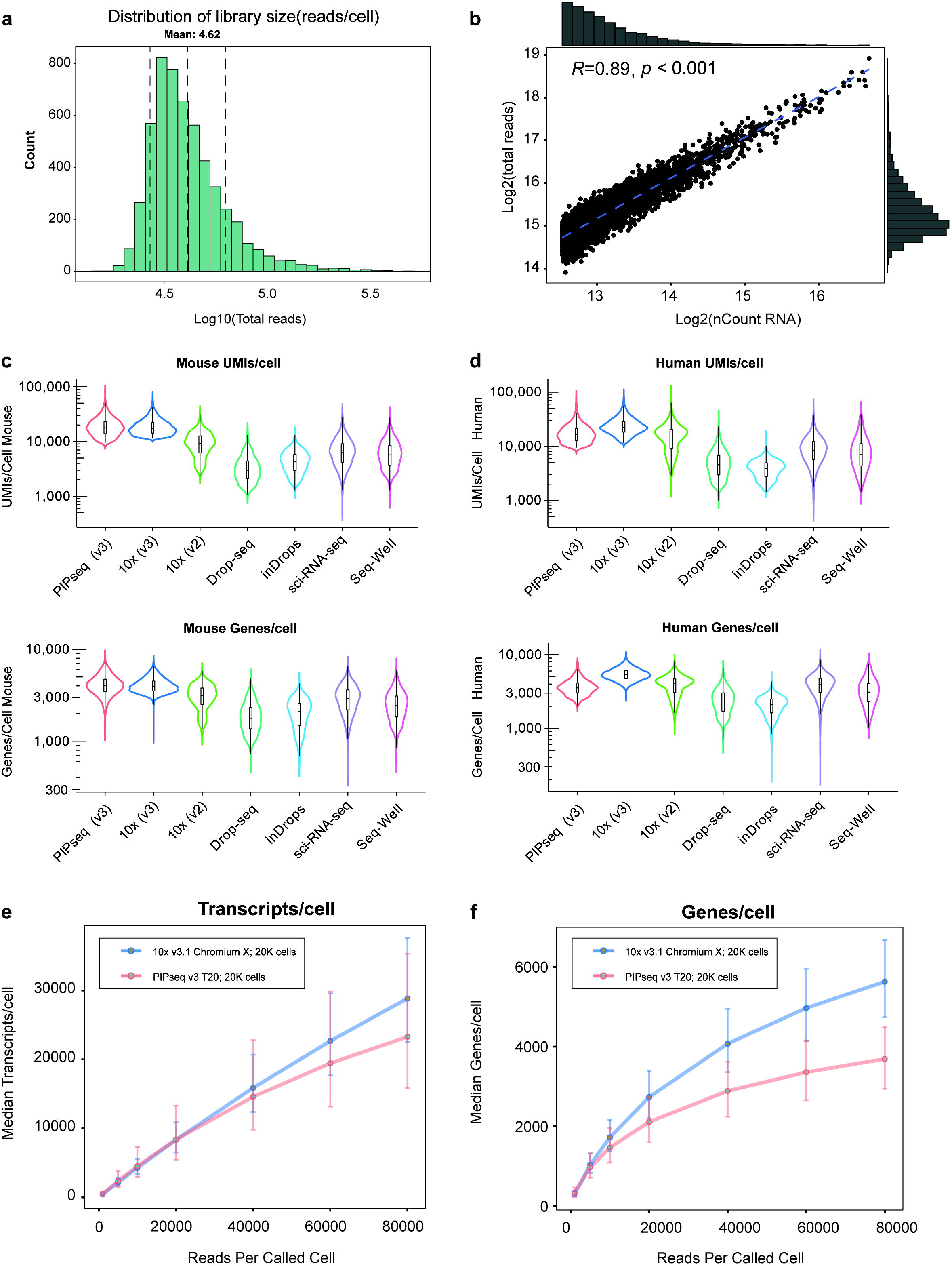
Data quality assessment of PIP-seq. **(a)** Representative distribution of reads per cell. **(b)** Correlation between reads and genes per cell. **(c,d)** Comparison of (c) UMIs/cell and (d) genes/cell in current single-cell methods. **(e,f)** Comparison of PIP-seq to 10X Genomics across a range of sequencing depths (0-80,000 reads/cell) (e) UMIs/cell and (f) genes/cell.

**Extended data Figure 6.**
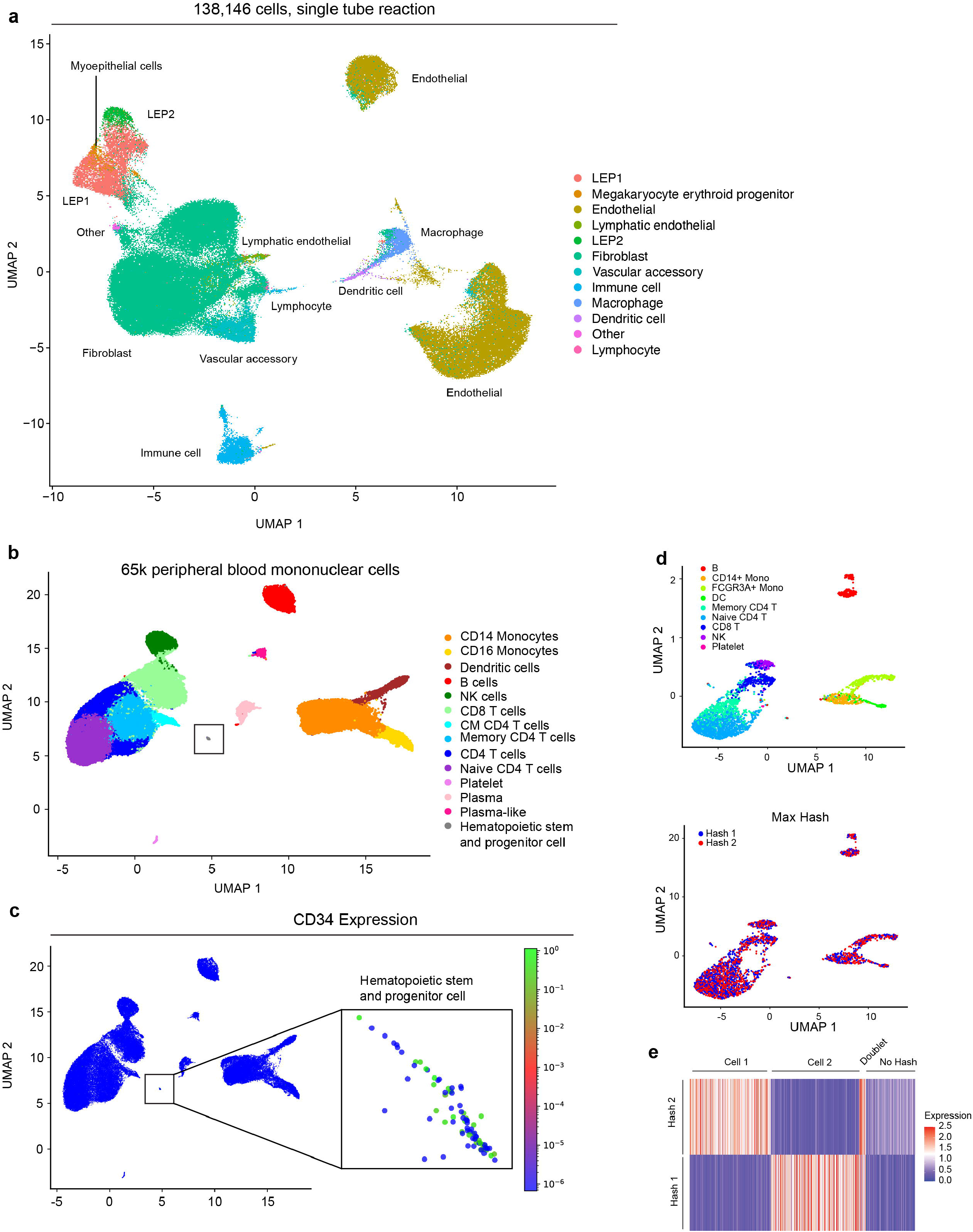
High cell number PIP-seq. **(a)** scRNA-seq of 138,146 single cells from breast tissue using a single-tube emulsification in 2-minutes. **(b)** scRNA-seq of 65k peripheral blood mononuclear cells (PBMCs) recovers a small population of CD34 cells. **(c)** Coloring of UMAP by the normalized expression of *CD34* for each cell. **(d)** Hashing of PBMCs demonstrates compatibility of PIP-seq with barcoded antibodies.

**Extended data Figure 7.**
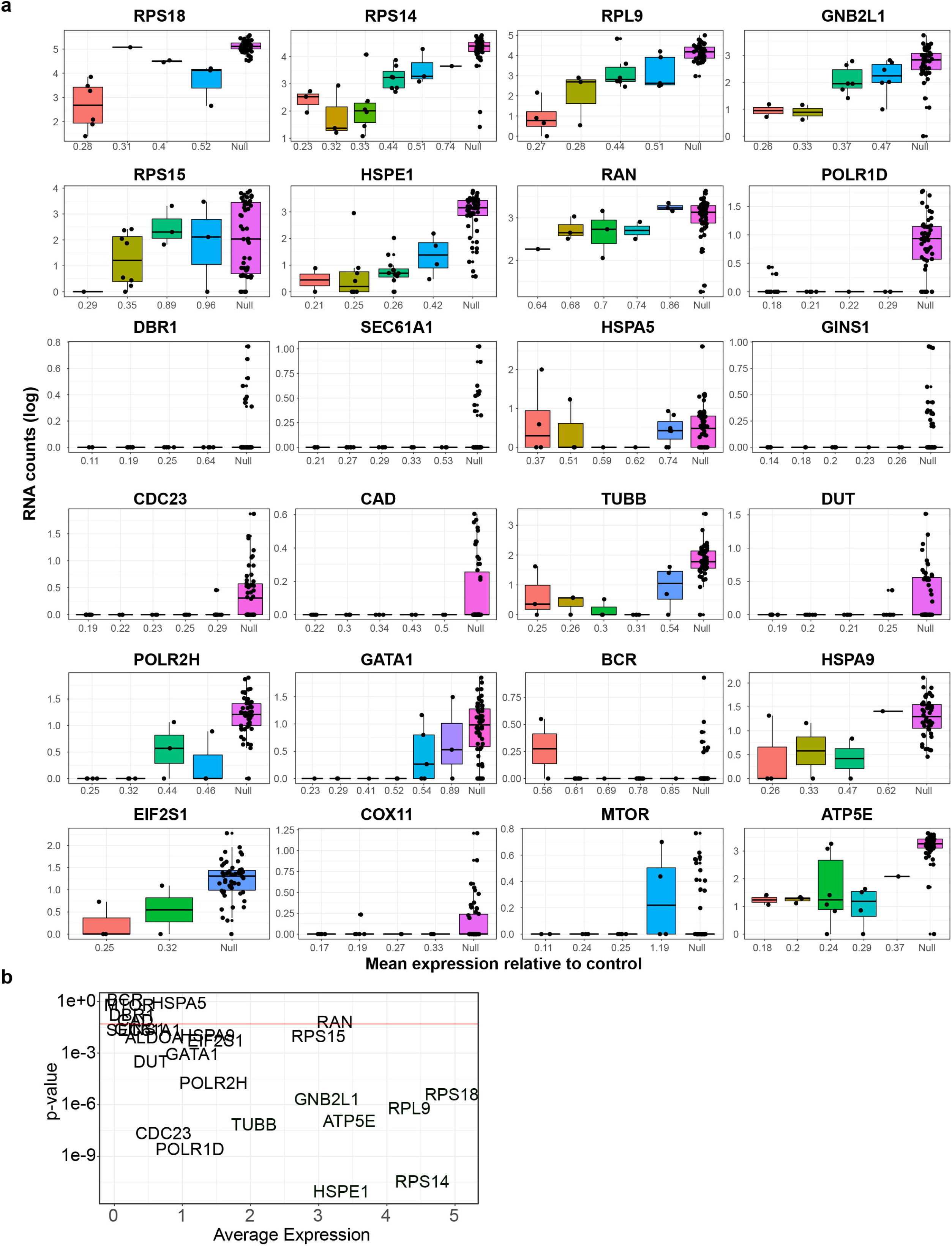
**(a)** Gene expression for each sgRNA within an allelic series for all genes in the CRISPRi library. Each sgRNA is ordered from predicted high to low knockdown efficiency. Non-targeting sgRNA are denoted as “Null.” **(b)** The relationship between gene expression and predicted knockdown of each gene. Expected changes in transcription across the allelic series were prominent in highly expressed genes. p-value represents the significance of the generalized additive model relating gRNA identity to knockdown efficiency for each gene.

**Extended data Figure 8.**
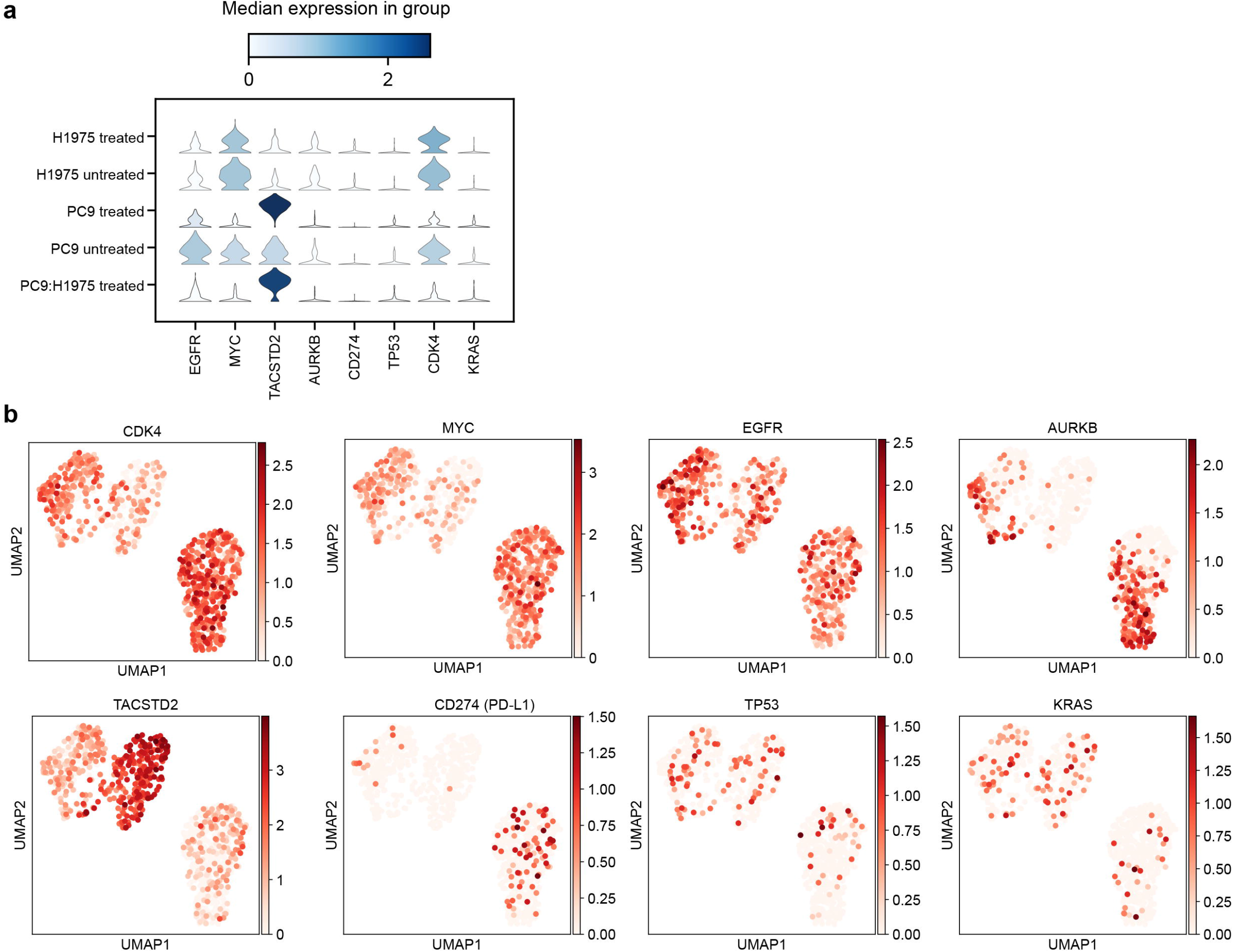
Identification of Gefitinib-specific transcriptional responses in cancer cell lines. **(a)** Violin plots of median expression values for selected differentially expressed genes. **(b)** The expression of selected differentially expressed genes superimposed on H1975 and PC9 cell clusters.

**Extended data Figure 9.**
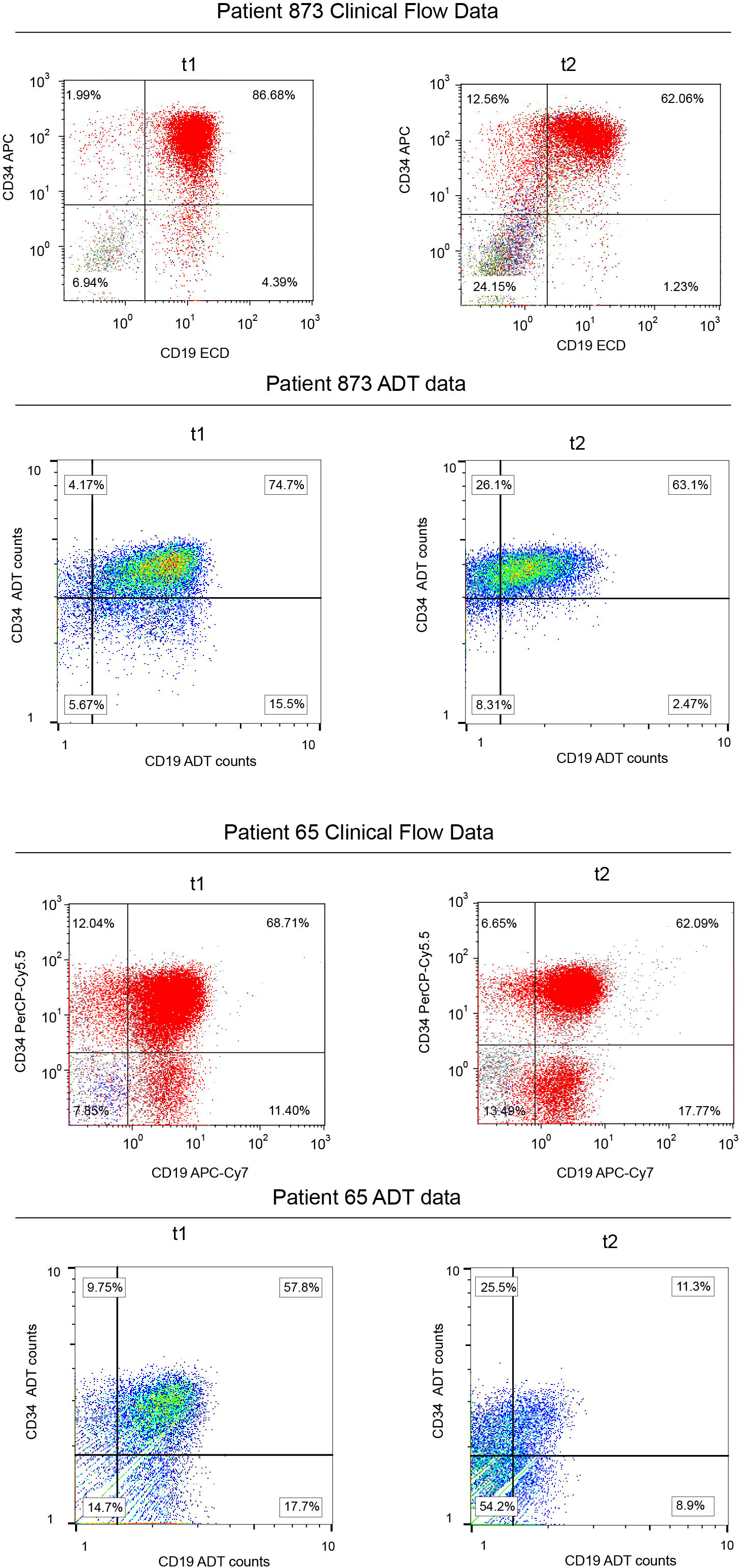
Clinical flow cytometry and corresponding antibody derived tag (ADT) data for patients 65 and 873 with mixed phenotypical acute leukemia (MPAL).

**Extended data Figure 10.**
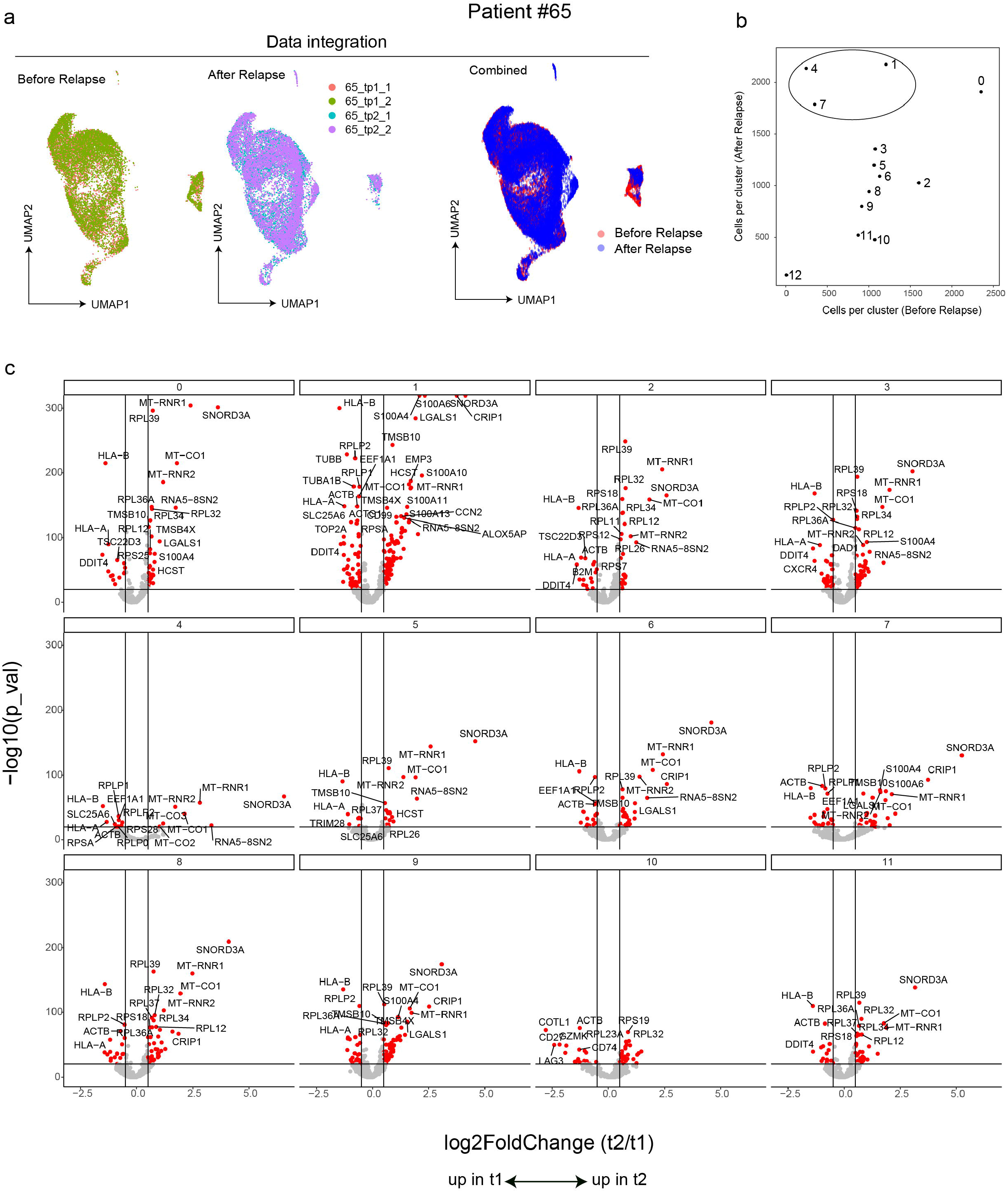
Analysis of PIP-seq data from MPAL Patient #65. **(a)** Integration of replicates and time points. **(b)** Correlation between the number of cells in each cluster before and after relapse identifies expansion of clusters 1,4, and7. **(c)** Volcano plots showing differential gene expression, by cluster, between t1 (before treatment) and t2 (after relapse).

**Extended data Figure 11.**
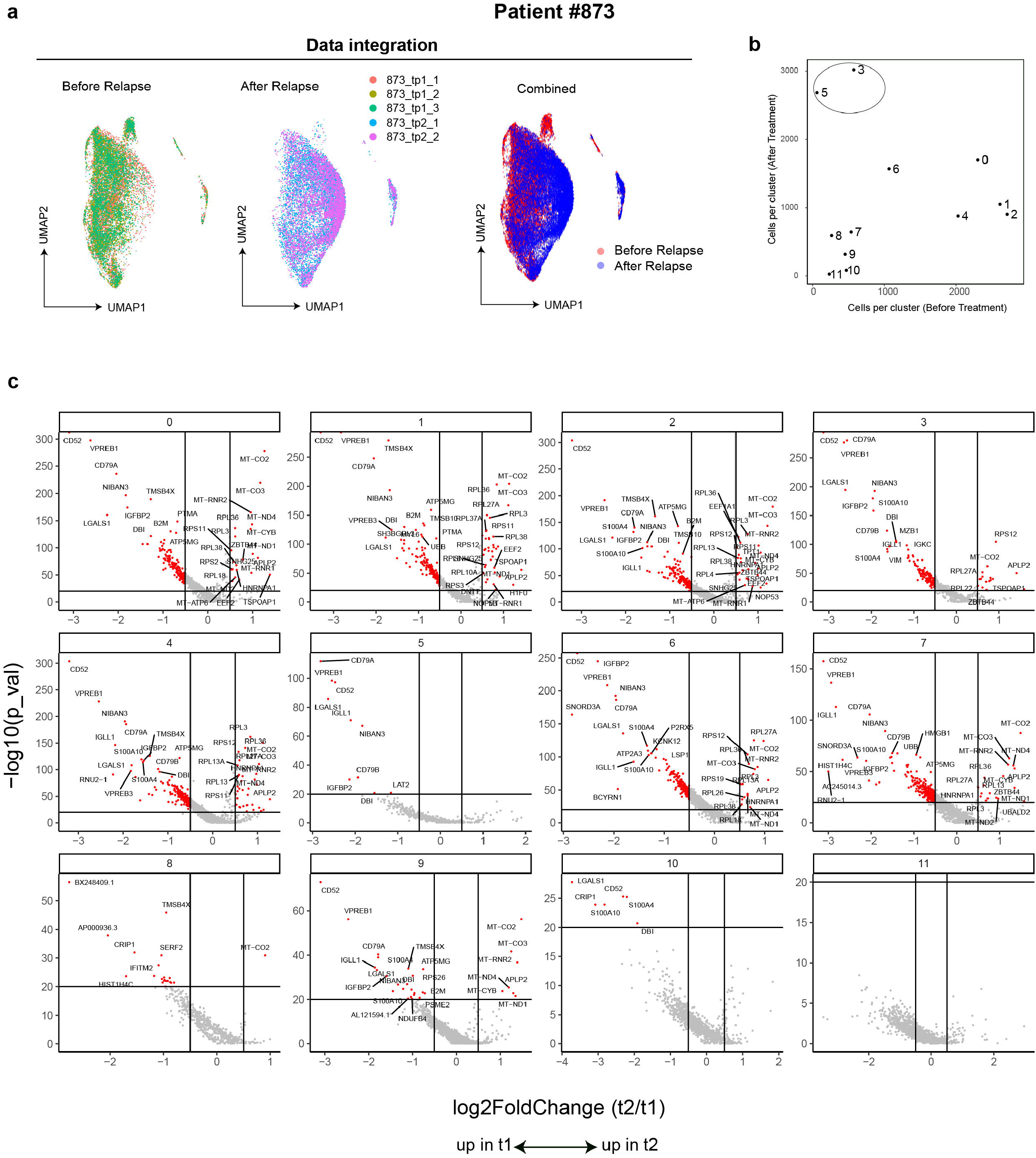
Analysis of PIP-seq data from MPAL Patient #873. **(a)** Integration of replicates and time points. **(b)** Correlation between the number of cells in each cluster before and after treatment identifies the expansion of clusters 3 and 5. **(c)** Volcano plots showing differential gene expression, by cluster, between t1 (before treatment) and t2 (after relapse).

## Tables

**Supplementary Table 1.** Oligonucleotides used in this study.

**Supplementary Table 2.** Breast tissue marker genes **(a)** used for cluster cell type identification and **(b)** identified using differential expression between clusters (FindAllMarkers Seurat v4.1.0, only.pos=FALSE, min.pct = 0.25, logfc.threshold = 0.5, test.use=‘bimod’).

**Supplementary Table 3.** Clinical characteristics of study participants.

**Supplementary Table 4.** Marker genes for patient 65 clusters identified using combined t1 and t2 timepoints (FindAllMarkers Seurat v4.1.0, only.pos=TRUE, min.pct = 0.25, logfc.threshold = 0, test.use=‘bimod’).

**Supplementary Table 5.** Marker genes for patient 873 clusters identified using combined t1 and t2 timepoints (FindAllMarkers Seurat v4.1.0, only.pos=TRUE, min.pct = 0.25, logfc.threshold = 0, test.use=‘bimod’).

**Supplementary Table 6.** Sequencing metrics.

**Supplementary Table 7.** PBMC marker genes (a) used for cluster-based cell type identification and (b) identified using differential expression between clusters.

## Acknowledgments

We thank members of the Abate laboratory for helpful advice and discussions.

## Author contributions

Conceptualization, ICC, ARA; Methodology, ICC, KMF, RHM, CD, ARA; Investigation, ICC, KMF, YX, DW, KTP, CDA, AO, JQZ, PH, JSAI, VK, CP; Formal Analysis, ICC, CH, AM-Z; Resources KMF, RHM, JMR, JSW, CCS, ZJG, ARA; Manuscript preparation, ICC, CCS, CACP, ARA; Supervision, ICC, KMF, RHM, CCS, ZJG, ARA.

## Data and materials availability

Sequencing data were deposited into GEO SuperSeries accession number GSE202919. The Reviewer access token is: ezipsugojfuvvql. All code will be made available from the corresponding author upon reasonable request.

