## Supplementary Table 3 for "Microfluidics-free single-cell genomics with templated emulsification"

| **Patient ID** | **Age** | **Gender** | **Timepoint** | **MPAL Subtype** | **Clinical Immunophenotype** | **Therapy** | **Best Response** |
| --- | --- | --- | --- | --- | --- | --- | --- |
| 65 | 54 | Male | Diagnosis | B/Myeloid | All blasts expressed CD34, CD38, HLA-DR, Majority subset: 82% of blast population expressed variable CD15, CD19, weak CD22, CD79a. Subset expressed TdT. Minority subset: 18% of blast population Expressed weak CD11c, weak CD13, CD33, CD64. | HDACi + Dauno | CR  (MRD by flow cytometry 1%) |
|  |  |  | Relapse | B/Myeloid | Expressed weak to absent CD15, CD19, CD38, Subset (69% of blast population) co-expressed CD34 and weak-to-absent CD22. |  |  |
| 873 | 76 | Female | Diagnosis | B/Myeloid | Expressed weak CD4, CD13, CD19, weak CD22, CD34, CD38, weak CD71, weak CD123, HLA-DR, Subset co-expressed CD117 (21% of gated events positive). Subset co-expressed CD10 (11% of gated events positive). | miniCVD + Ino | None  (MRD by flow cytometry 70%) |
|  |  |  | Relapse | B/Myeloid | Expressed weak CD4, CD13, variable CD19, weak-absent CD22, CD33, bright CD34, weak CD38, weak-absent CD64, weak CD71, variable CD117, weak CD123, weak HLA-DR, Minor subset expressed possible CD10. |  |  |

CR = Complete Response; HDACi = histone deacetylase inhibitor; Dauno = daunorubicin; MiniCVD = cyclophosphamide and dexamethasone at 50% dose reduction. Ino.= inotuxumab ozogamicin; MRD = measurable residual disease
